# A small-scale shRNA screen in primary mouse macrophages identifies a role for the Rab GTPase Rab1b in controlling *Salmonella* Typhi growth

**DOI:** 10.1101/2020.12.01.406090

**Authors:** Virtu Solano-Collado, Rosa Angela Colamarino, David A Calderwood, Massimiliano Baldassarre, Stefania Spanò

**Affiliations:** Institute of Medical Sciences, University of Aberdeen, Foresterhill, Aberdeen AB252ZD, United Kingdom; Department of Pharmacology, Yale University School of Medicine, New Haven, CT, USA; Department of Cell Biology, Yale University School of Medicine, New Haven, CT, USA

**Keywords:** Host-defence, *Salmonella* Typhi, Rab GTPases, Rab1b, shRNA screen, macrophages, RAB GTPases

## Abstract

*Salmonella* Typhi, the causative agent of typhoid fever, a life-threatening systemic infection, is a human restricted bacterial pathogen. A fundamental aspect of *S*. Typhi pathogenesis is its ability to survive in human macrophages but not in macrophages from other animals (i.e. mice). Despite the importance of macrophages in establishing systemic *S*. Typhi infection, the mechanisms that macrophages use to control the growth of *S*. Typhi and the role of these mechanisms in the bacterium’s adaptation to the human host are mostly unknown. To facilitate unbiased identification of genes involved in controlling the growth of *S*. Typhi in macrophages, we report optimized experimental conditions required to perform loss-of function pooled shRNA screens in primary mouse bone-marrow derived macrophages. Following infection with a fluorescently-labeled S. Typhi, cells defective in genes important for controlling *S*. Typhi growth (and therefore unable to kill *S.* Typhi) are sorted based on the intensity of fluorescence (i.e. number of intracellular fluorescent bacteria). shRNAs enriched in the fluorescent population are identified by next-generation sequencing. A proof-of-concept screen targeting the mouse Rab GTPases confirmed Rab32 as important to restrict *S*. Typhi in mouse macrophages. Interestingly and rather unexpectedly, this screen also revealed that Rab1b controls *S*. Typhi growth in mouse macrophages. This constitutes the first report of a Rab GTPase other than Rab32 involved in *S*. Typhi host-restriction. The methodology described here should allow genome-wide screening to identify mechanisms controlling the growth of *S*. Typhi and other intracellular pathogens in primary immune cells.

## Introduction

Upon bacterial invasion, cells of the innate immune system such as macrophages act as the first line of defense to control the infection. Macrophages are equipped with an array of antimicrobial factors, allowing them to efficiently eliminate invaders (Weiss and Schaible, 2015). However, some bacterial pathogens have evolved mechanisms to resist the attack by macrophages and adopt an intracellular lifestyle, persisting inside these cells. The Gram-negative pathogen *Salmonella enterica*, upon internalization by a host cell, establishes an intracellular niche by creating a specialized compartment known as *Salmonella*-containing vacuole where the bacterium survives and replicates. The capability of *Salmonella* to exploit and survive within macrophages is essential for its pathogenesis and for the establishment of systemic infection (Fields et al., 1986). This intracellular survival is made possible by the action of type-III secretion systems (T3SS) that inject effector proteins into target cells to manipulate host pathways. *Salmonella* encodes two T3SS within the pathogenicity islands 1 (SPI-1) and 2 (SPI-2). Some of these effectors block specific macrophage killing mechanisms while others facilitate bacterial invasion and create a habitable intracellular niche (Jennings et al., 2017, Galan et al., 2014, Spano et al., 2016).

The species *Salmonella enterica* comprises several serovars or serotypes (based on their surface antigen composition) and while most can infect a broad range of mammalian species others are host specific or host-restricted. *Salmonella enterica* serovar Typhi (*S*. Typhi) is a host-restricted serovar that only infects humans (Dougan and Baker, 2014) and causes the life-threatening condition typhoid fever, resulting in between 11.9 and 26.9 million cases and hundreds of thousands of deaths annually around the globe (Gibani et al., 2018). The failure of this pathogen to infect other species is partially due to its inability to overcome a defence mechanism, initially identified in mouse macrophages, which depends on the Rab GTPase Rab32 and its nucleotide exchange factor Biogenesis of Lysosome-related Organelles Complex-3 (BLOC-3) (Spano et al., 2016, Spano and Galan., 2012). This pathway is efficiently disarmed by the broad-host serovar *S*. Typhimurium through the delivery of two T3SS effectors: GtgE and SopD2. Both act directly on Rab32, GtgE is a protease and a SopD2 a GTPase activating protein (Spano et al., 2016).

Rab GTPases are key regulators of vesicle trafficking between organelles and an interface between the host cell and the internalized pathogen. Thus, it is not surprising that successful intracellular pathogens use the manipulation of these proteins as a mechanism to survive within their eukaryotic hosts (Stein et al., 2012). *Mycobacterium tuberculosis*, *Legionella pneumophila*, *Coxiella burnetii* and *S.* Typhimurium are examples of well-studied intracellular pathogens that actively modulate Rab proteins to establish an intracellular niche and subvert macrophage killing (McGourty et al., 2012, Bakowski et al., 2010, Spano and Galan, 2018, Hardiman et al., 2012, Seto et al., 2011). Despite efforts to understand the nature of *S*. Typhi host-restriction, most of our knowledge of *Salmonella* interactions with Rab GTPases is extrapolated from *S*. Typhimurium studies, and many questions as to how macrophages deal with this important human pathogen remain. In this study, using a loss-of-function shRNA-based approach, we have performed a screen in mouse bone-marrow derived macrophages and identified Rab1b as factor controlling *S*. Typhi growth in mouse macrophages.

## Materials and Methods

### Bacterial strains and mammalian cell culture

The *S*. Typhi wild-type ISP2825 (Galan and Curtiss., 1991) and the *S*. Typhi *glmS::Cm::mCherry* (*S*. Typhi::*mCherry*) (Baldassarre et al., 2020) strains have been described previously.

*Salmonella* can induce caspase 1-dependent macrophage death (Brennan and Cookson, 2000), which could complicate the interpretation of the results. To avoid this, bone-marrow-derived macrophages (BMDMs) were isolated from caspase 1-deficient macrophages (casp1_−/−_) and differentiated as described before (Weischenfeldt and Porse., 2008). Briefly, BMDMs were collected from mouse femurs and differentiated in RPMI 1640 medium (Thermo Fisher) supplemented with 10% fetal bovine serum (FBS; HyClone), 2 mM glutamine (Thermo Fisher), 10 mM Hepes (Thermo Fisher), 1 mM Sodium Pyruvate (Thermo Fisher) and 20% L929 cells supernatant as the source of macrophage stimulating factor (M-CSF).

Immortalised bone-marrow-derived macrophages (iBMDMs), RAW264.7 and HEK293T cell lines were routinely maintained in DMEM high glucose 2mM glutamax (Thermo Fisher) and 10% FBS.

### Infection of macrophages with *Salmonella* Typhi for intracellular survival assays (CFU assay)

Overnight cultures of *Salmonella* were diluted 1/20 in LB broth containing 0.3 M NaCl and grown at 37°C in a rotary wheel for 2 hour and 45 minutes. Cells were washed twice with Hank’s Balanced Salt Solution (HBSS, Thermo Fisher) and infected with the different strains of *S*. Typhi at the indicated multiplicity of infection. One hour post-infection, cells were washed twice with HBSS and incubated in growth medium supplemented with 100 μg/ml gentamicin for 30 min to kill extracellular bacteria. Then, medium was replaced with growth medium containing 5 μg/ml of gentamicin to avoid cycles of reinfection. At the indicated times-post infection, cells were washed twice with phosphate buffer saline (PBS, Sigma), lysed in 1 ml 0.1% sodium deoxycholate (Sigma) in PBS and intracellular bacteria counted by plating serial dilutions on LB-agar plates.

### *Salmonella* Typhi infection of macrophages for flow-cytometry and fluorescence microscopy

To determine the optimal multiplicity of infection (MOI), 6×10^5^ BMDMs were plated on 6-well plates and infected with different amount of fluorescent bacteria (from moi 5 to 30). One hour-post infection, medium was replaced with growth medium containing 100 μg/ml gentamicin to kill extracellular bacteria. Cells were then collected from the plate by incubating with cold Versene for 5 minutes, fixed with 4% paraformaldehyde (PFA) for 10 minutes at room temperature and then analysed using the LSRFortessa flow cytometer. In parallel, 1×10^5^ BMDMs were plated on 24-well plates and infected as before. Cells were then fixed with 4% paraformaldehyde (PFA) for 10 minutes at room temperature and visualised with a Zeiss Imager M1 fluorescence microscope. For quantification of the number of bacteria per cell, 85 (out of 272; moi 5), 83 (out of 190; moi 10), 72 (out of 147; moi 15), 80 (out of 128; moi 20) and 88 (out of 122; moi 30) infected cells were analyzed and the number of bacteria per cell plotted with Prism 8.

To confirm that shRNA-mediated knockdown resulted in a measurable phenotype in BMDMs, 4×10^6^ BMDMs were transduced with shRNAs targeting HPS-1 or scrambled sequence and plated on 6-well plates (6×10^5^ cells/well; flow cytometry) or 24-well plates (1×10^5^ cells/well; fluorescence microscopy and CFU assays). After selection with puromycin, transduced cells were infected with *S*. Typhi::*mCherry* and samples at 1.5, 5 and 24 hours post-infection were analysed using the LSRFortessa flow cytometer and fluorescence microscopy as described above. For quantification of the number of bacteria per cell, a minimum of 100 cells were used per condition.

### RNA isolation and RT-qPCR

Total RNA from eukaryotic cells was isolated using the RNeasy mini kit (QIAGEN). RNA was transcribed using the iScript reverse transcriptase (BioRad) and transcript levels were determined using the Takyon SYBR MasterMix (Eurogentec) and the StepOnePlus real-time PCR system (Applied BioSciences).

### Generation of Rab GTPases shRNA library and screening

The mouse Rab GTPases shRNA library (317 shRNAs) was extracted from The Mission mouse shRNA library generated by The RNAi consortium (Sigma Aldrich) and prepared as described before (Shi et al., 2016). The shRNA targeting the HPS-1 gene (TRCN0000292556) as well as the packaging plasmids pCMV-VSV-G and pCMV-dR8.91 were obtained from Sigma Aldrich. The plasmids encoding the shRNA sequences were pooled in equal concentrations and mixed with the packaging plasmids pCMV-VSV-G and pCMV-dR8.91 to generate lentiviral particles. HEK293T cells were transfected with the plasmid mixture and polyethyleneimine (PEI) as transfecting reagent. Fifty-two hours post-transfection, the lentivirus particles were collected from the cell supernatant.

For each screen, a total of 10×10^6^ BMDMs were transduced at day 4 after isolation using a dilution 1:10 of the lentiviral pool for 24 hours and selected in puromycin (5 μg/ml) for 48 hours. Cells were then infected with S. Typhi::*mCherry* at an MOI of 10. At 1.5 and 24 hours-post infection, cells were lifted from the plate by incubating with cold Versene for 5 minutes and fixed with 4% PFA for 10 min at room temperature. Three independent experiments were performed and sorted independently. A total of 50,000 cells (non-infected and with low number of intracellular bacteria) or 17,000 to 20,000 cells (high number of intracellular bacteria) were sorted in a BD Influx BSLII Sorter (Ian Fraser Cytometry Centre, University of Aberdeen). For genomic DNA isolation, the DNeasy Blood & Tissue kit (QIAGEN) was used following the instructions of the manufacturer with the following modifications: after cell lysis, samples were incubated at 90°C for 1 hour to reverse PFA modification of DNA. The resulting gDNA was used as template for PCR amplification of the shRNA encoding regions using the Fw-NGS and the Rev-NGS primers (IDT Technologies) and the Phusion High Fidelity DNA polymerase (NEB). Three PCRs per condition were performed and the amplicons combined. Primer sequences are listed in Table S5. A second PCR was performed to attach indexes and Illumina adaptors and the PCR amplicon was sequenced on the NextSeq500 v2 instrument (Centre for Genome-Enabled Biology and Medicine (CGEBM), University of Aberdeen).

### Data analysis

The analysis of the data involved background correction and scoring. Firstly, the proportion of reads relative to total reads of each shRNA was obtained using the BBDUK software (CGEBM, University of Aberdeen). shRNAs with value “0” in 2 out of the 3 experiments in the plasmid library or controls (non-sorted cells) at either 1.5 and 24 hours-post infection were excluded from the analysis (see Table S2). Then, the proportion of reads of each shRNA within the high fluorescence population (H) relative to the total proportion of reads (H, M and N populations) was calculated as well as the average and standard deviation of the population. Finally, the z-score for each shRNA was calculated as z = (x-μ)/σ, where x is the value obtained of each shRNA (proportion of reads of each shRNA within the high fluorescence population (H) relative to the total reads (N+M+H)), μ is the population mean and σ is the standard deviation of the population (see Table S3). True hits were defined as those for which at least 2 shRNAs had a z-score ≥1.18 based on the scores obtained for internal positive controls. For each gene, the second best shRNA was validated (see Table S4).

### Validation of candidate genes by individual gene knockdown and intracellular survival assay

iBMDMs were transduced with lentivirus encoding shRNA to knockdown individually candidate genes 6 days before infection. Twenty-four hours after transduction, cells were incubated with 5 μg/ml puromycin for 2-4 days. *S*. Typhi infection assays were carried out as described before.

## Results

### shRNA-mediated knockdown in bone-marrow derived macrophages (BMDMs)

Pooled shRNA screens offer unbiased approaches to identify genes important for specific of biological processes (Carette et al., 2011, Silva et al., 2008). As illustrated in Figure 1, we designed an shRNA screening strategy to identify host macrophage proteins whose knockdown facilitated *S*. Typhi growth. The approach requires efficient delivery of the shRNA library into primary macrophages to generate a pool of knockdown cells coupled with an effective sorting strategy to collect cells that failed to clear the invading bacteria. As described below we optimized each of these steps.

**Figure 1.**
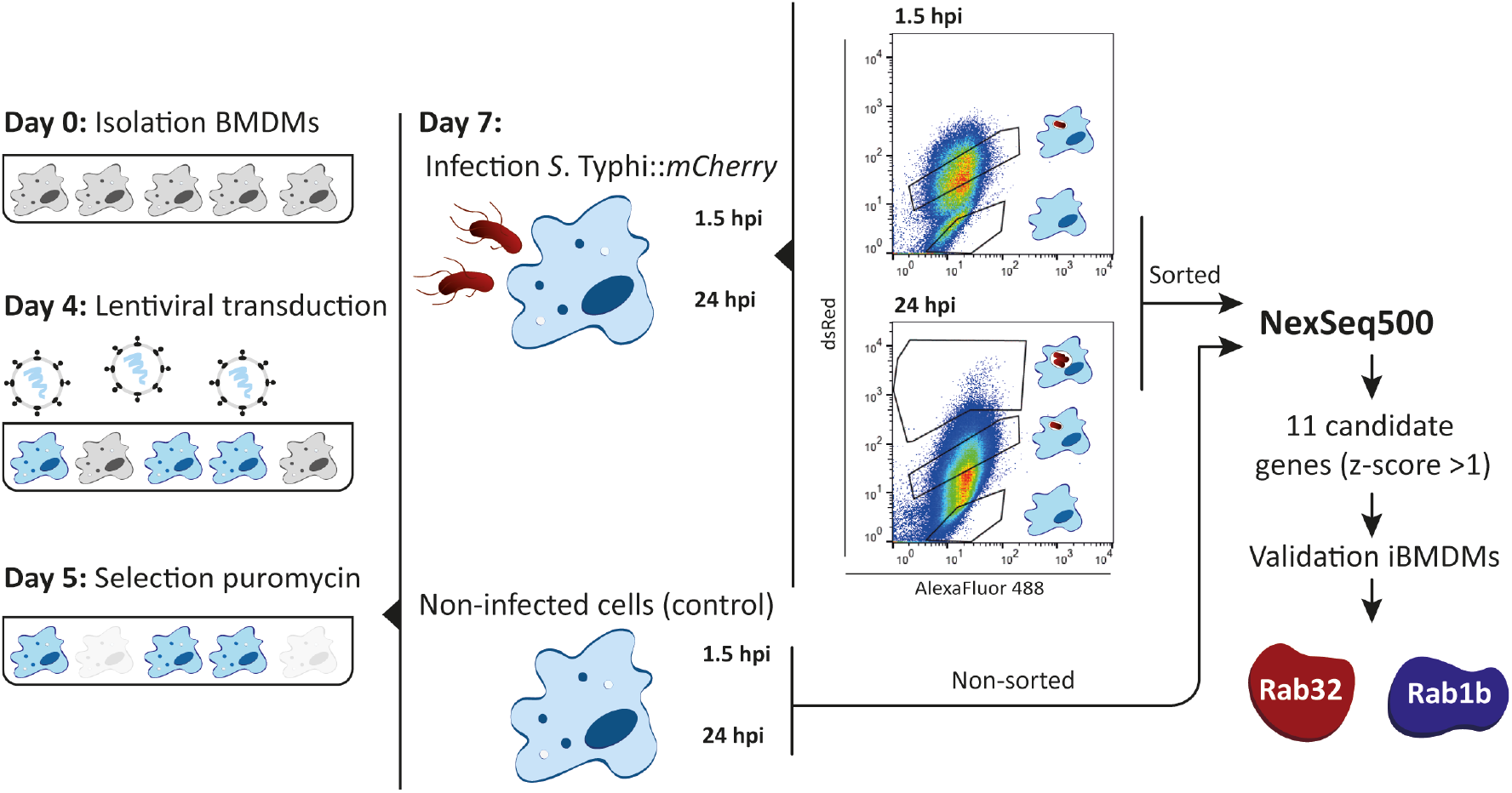
shRNA screening in BMDMs. Schematic representation of the experimental approach of the shRNA screening in BMDMs. BMDMs were isolated from casp1−/− mice and transduced at day 4 after isolation. After selection with puromycin for 48 hours, cells were infected with S. Typhi::*mCherry*. At 1.5 and 24 hours post-infection cells were fixed with 4% PFA and the different sub-populations sorted based on dsRed fluorescence (gates for the different sub-populations indicated). Genomic DNA from sorted and non-sorted cells (control) was isolated, the region encoding the shRNA amplified by PCR and the amplicons sequenced by next-generation sequencing (NextSeq500). As a result of the screen, 11 candidate genes were identified based on a z-score >1.18 and hits validated using shRNAs to knockdown the individual genes.

Macrophages are difficult to manipulate genetically, probably because they are terminally differentiated. Therefore, a requisite for large-scale loss of function screening in primary macrophages is optimization of conditions to achieve efficient and reproducible gene knockdown within the macrophage lifespan and without affecting their final differentiation. Although several methods to introduce exogenous DNA in primary murine macrophages have been reported (Thompson et al., 1999, Guiet et al., 2012), lentiviral systems have been shown to be the most efficient method (Naldini et al., 1996, Schroers et al., 2000). While Zeng et al., have shown that culturing macrophages for 8 days *in vitro* increased the transduction efficiency using HIV-based vectors (Zeng et al., 2006) others recommend earlier stages of differentiation for efficient transduction (Miller and Blystone., 2015). These differences might be due to the methods and/or the vectors used to generate the lentiviral particles and/or the methods used to differentiate BMDMs. In order to determine, under our differentiation conditions, and with our lentiviral system, the optimal post-isolation day to transduce BMDMs we used a lentivirus encoding an shRNA targeting the Rab GTPase Rab32 and the puromycin resistance marker. We transduced BMDMs that were in culture from day 4 to day 9 of differentiation. Transduced cells were selected with 5 μg/ml puromycin for 48 hours and cell survival determined by alamar blue assays. The results obtained showed that at the early stages of the differentiation into macrophages (day 4), cells are transduced more efficiently than at later stages (days 7 to 9) (Figure 2A). Importantly, the level of transduction achieved (≈60%) ensured that the probability of having a cell infected by more than one virus (and then more than one shRNA) is low.

**Figure 2.**
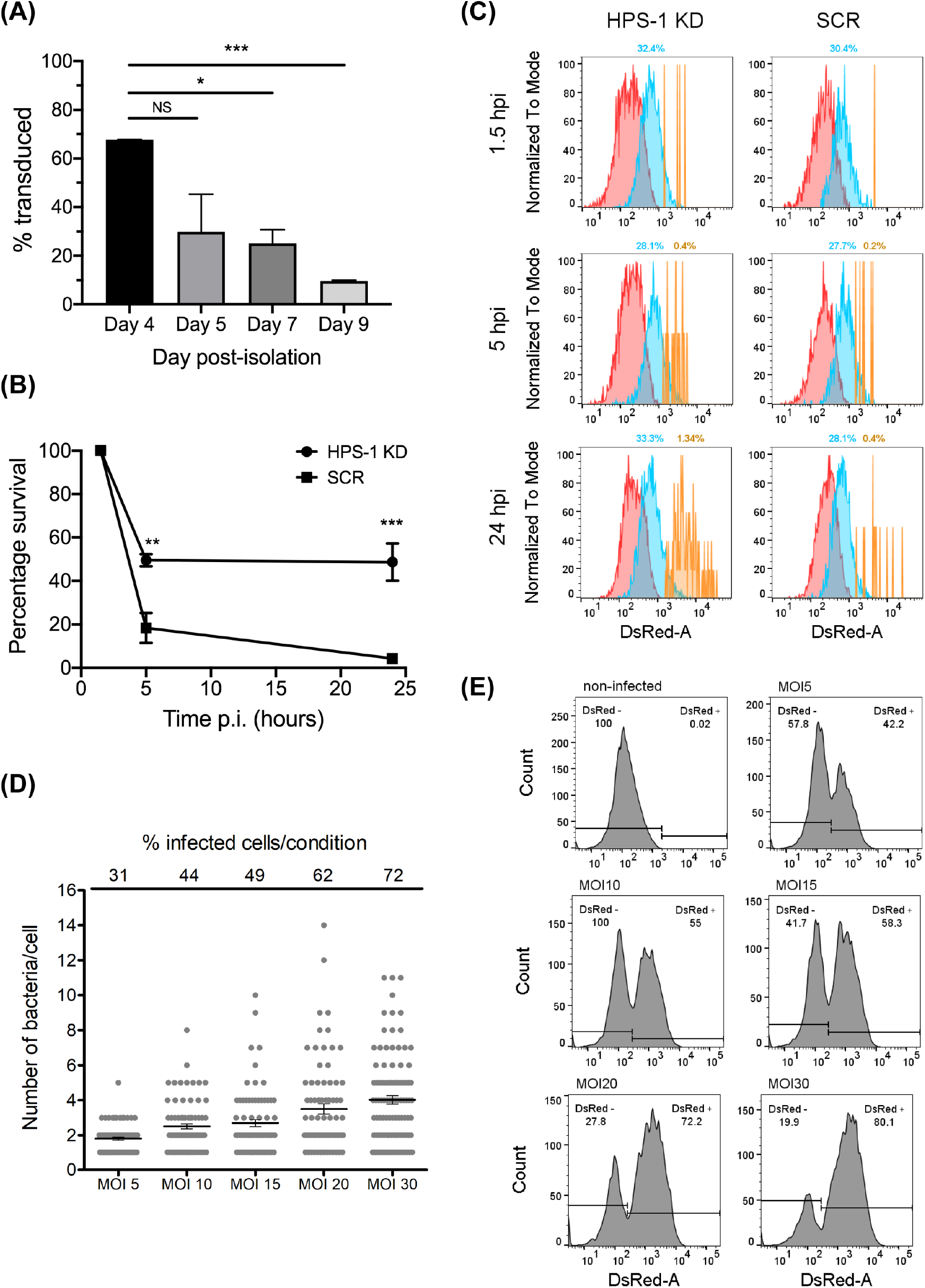
shRNA-mediated knockdown in bone-marrow derived macrophages (BMDMs). **(A)** BMDMs were transduced at different days post-isolation. After incubation with 5 μg/ml puromycin for 48 hours, cell viability was evaluated with alamar blue assay. Percentage of survival are: Day 4 67.72%, Day5 26.84%, Day 7 25.02%, Day 9 9.63%. Values are mean ± SEM of two independent measurements. NS=non-significant; \**P<0.05*; \*\*\**P<0.005*. **(B)** BMDM cells were transduced with lentiviral particles encoding a non-target scrambled sequence (SCR) or encoding a sequence targeting HPS-1 (HPS-1 KD). After selection with 5 μg/ml puromycin, cells were infected with *S*. Typhi wild-type, lysed at three times post-infection (1.5, 5 or 24 h) and CFUs were enumerated. Values are means ± SEM of two independent experiments performed in triplicate. *P* values were determined by the Student’s *t* test and *P* values indicated when *P*<0.05. **(C)** BMDMs HPS-1 Knockdown (HPS1 KD) or control (Scrambled shRNA; S) were infected with *S*. Typhi::*mCherry* (MOI 10). At different times post-infection (1.5, 5 and 24 hours post-infection), cells were fixed with 4% PFA and analyzed by flow cytometry. The three subpopulations non-infected (N) (red), 1-3 bacteria (M) (blue) and >3 bacteria (H) (yellow) are shown and the percentage of the M and H populations are indicated. **(D-E)** BMDMs were infected with *S*. Typhi::mCherry at different multiplicity of infection (MOI). After 1.5 hours-post infection, cells were fixed with 4% PFA and analyzed by fluorescence microscopy **(D)** or flow cytometry **(E)** to determine the average number of bacteria per cell. The percentage of infected cells per condition is indicated.

To confirm that the viral transduction process did not affect the ability of mouse macrophages to control *S*. Typhi growth, we transduced BMDMs with a virus encoding a non-targeting shRNA scrambled sequence (SCR) and assessed *S*. Typhi intracellular survival by CFU assays (Figure 2B). The results obtained confirmed that viral-transduced BMDMs are able to control *S*. Typhi growth. Next, to confirm that shRNA gene knockdown results in a measurable phenotype in BMDMs, we knocked-down HPS-1 (≈95% knockdown; Figure S1). HPS-1 is one of the two subunits of the BLOC-3 complex, and BLOC-3 together with Rab32, controls *S*. Typhi growth in macrophages (Spano and Galan., 2012, Baldassarre et al., 2020). We initially assessed *S*. Typhi intracellular survival in HPS-1-depleted macrophages using CFU assays (Figure 2B). The results obtained were further confirmed by flow cytometry and fluorescence microscopy (Figure 2C and S2). For this, HPS-1 knockdown BMDMs were infected with *S*. Typhi*::mCherry* and samples were analyzed at three different times post-infection: 1.5 (internalized bacteria), 5 and 24 hours. As shown in Figure 2C, the analysis by flow cytometry showed a population representing 1.34% of the HPS-1 depleted cells (≈4% of infected cells) containing higher loads of intracellular bacteria at 24 hours post-infection, while this population only represents 0.4% of control cells. Similarly, the analysis by fluorescence microscopy showed that the number of intracellular bacteria per cell significantly increased at late time points in HPS-1 knockdown cells but not in control cells. (Figure S2).

Taken altogether, these data confirm that viral-delivered shRNAs per se, do not affect the macrophages general behavior and that shRNA-dependent gene knockdown results in an assessable phenotype in BMDMs.

### Flow cytometric analysis of *S*. Typhi infection in BMDMs

Our screening approach (Figure 1) relies on an efficient flow cytometric method to rapidly detect and isolate cells that, upon gene silencing, are more permissive to *S*. Typhi growth (i.e. contain high numbers of intracellular *S*. Typhi). However, if a macrophage lacks a factor required to control the growth of *S*. Typhi, replication of the bacteria within the cell could ultimately lead to cell death and loss of a positive hit. To limit this we have optimized the ratio of bacteria/cell during the infection (multiplicity of infection; MOI) in order to obtain a reasonable infection rate (50-60% of infected cells) with low numbers of bacteria in each infected cell (2-3 bacteria per cell). For this, BMDMs were infected with different amounts of *S*. Typhi*::mCherry*, from MOI 5 to 30. One-hour post infection, cells were treated with gentamicin for 30 minutes to kill any extracellular bacteria. Cells were then fixed and the percentage of infected cells, as well as the number of bacteria per cell, was determined by fluorescence microscopy (Figure 2D) and flow cytometry (Figure 2E). When cells were infected with an MOI of 5, the mean number of intracellular bacteria was 2 and the percentage of infected cells was around 30% (42.2% flow cytometry). In cells infected with an MOI of 10, the percentage of infected cells increased to 44% (55% flow cytometry) and only 9 out of 83 infected cells contained more than 4 intracellular bacteria. Higher MOI resulted in higher number of infected cells (from 42.2% to 80.1%) but the number of bacteria per cell increased notably as well, finding a considerable number of cells with more than 6 bacteria per cell with the highest MOI (Figure 2D). Therefore, all subsequent experiments were performed using an MOI of 10. In addition, these analyses allowed us to determine the sensitivity of the flow cytometer. Cells with as low as 1-2 fluorescent intracellular bacteria were detectable as an independent cellular population and easily distinguishable from non-infected cells (Figure 2E).

### shRNA screen of Rab GTPases involved in *S.* Typhi growth restriction in mouse macrophages

In order validate our screening strategy and to identify novel Rab GTPases involved in restricting the growth of the human pathogen *S*. Typhi, we performed a targeted loss-of-function shRNA screen in BMDMs using a custom-made pooled library targeting the mouse Rab GTPases (Table S1; summarized in Figure 1). In addition, we included a validated shRNA targeting HPS-1 as an internal positive control for the screen.

Three independent screens were performed. For each one 10×10^6^ BMDMs were transduced at 4 day after isolation with a pool of lentiviruses encoding the library and the positive control (see Materials and Methods). After 48 hours of selection with 5 μg/ml puromycin the antibiotic was removed and the cells were infected with *S*. Typhi*::mCherry* at MOI 10. One hour after infection, the medium was replaced with fresh medium containing 100 μg/ml gentamicin for 30 min to kill extracellular bacteria. At 1.5- and 24-hours post-infection, cells were lifted from the plates, fixed with 4% PFA and the different sub-populations of cells sorted based on red-fluorescence (an example is shown in Figure S3). Based on our fluorescence experiments, the sorted populations contains on average no bacteria (N), 1-3 bacteria (M) >3 bacteria (H).

This sorting strategy enabled enrichment of the very low abundant population containing more than 3 bacteria (from 0.15-0.2% of the total number of cells up to 0.5%). Once sorted, the genomic DNA from each cellular sub-population was isolated and regions containing the shRNA sequences were PCR-amplified (3 PCRs per sample per experiment), the replica for each experiment pooled and the amplicons subjected to next-generation sequencing. Amplicons obtained from both the plasmid library and non-sorted cells were sequenced and used as controls to determine i) the abundance of each shRNA in the initial plasmid pool (plasmid library), ii) whether a particular hairpin present in the plasmid library was lost after transduction (absent from the non-sorted 1.5 hours post-infection sample) and iii) whether a hairpin was lost over time independently of the infection (absent from the non-sorted 24 hours post-infection sample) (see Figure 1 and Table S2).

Out of the 318 shRNAs that comprised the library, 7 shRNAs were not present or not detected by deep sequencing (0 reads in the “plasmid library” sample), 8 shRNAs were not present in non-sorted samples at either 1.5 or 24 hours and the reads for 2 shRNAs were “0” in all the sub-populations in at least one full experiment. All those shRNAs were not considered for further analyses (see Table S2). Importantly, using these criteria none of the 56 genes included in the library were lost and at least 3 shRNAs per gene were recovered and included in the analysis. To obtain the list of candidate genes involved in *S*. Typhi growth-restriction and therefore, those more abundant in the H population (see Figure 1 and S3), we first calculated for each shRNA the total proportion of reads (N, M and H) as well as the proportion of reads within the high fluorescent sub-population (H). Then, we calculated the percentage of reads in H relative to total reads (N, M and H). Finally, we calculated the z-score for each individual shRNA as z = (x−μ)/σ, where x is the final value obtained of each shRNA, μ is the population mean and σ is the standard deviation of the population (Figure S3B and Table S3). By performing this analysis, the z-score values obtained for the positive controls were: HPS-1, z = 1.22 and Rab32 z = 1.18, z = 2.29, z = 0.63, z = −0.37 and z = −1.18). In view of the scores obtained for known positive hits, we decided that a less stringent strategy would be preferable in order to minimize the chances of excluding positive candidates. Therefore, we selected 1.18 as threshold to call out hits and used the following criterion to generate a candidate gene list: 2 or more shRNAs with a z-score >1.18. Using this strategy we identified 11 candidate genes (≈20 % of the total genes) that are listed in Table S4.

### Rab1b is important to control *S*. Typhi growth in mouse macrophages

To prove the reproducibility of the data independently of the cell model, we used immortalized BMDMs (iBMDMs) to validate our hits. Additionally, this allowed us to reduce the use of animals in line with the principles of the 3Rs (Replace, Reduce, Refine). We first confirmed that iBMDMs behave similarly to BMDMs in their ability to kill *S*. Typhi (Figure 3A). We also confirmed that HPS-1 depleted iBMDMs are more permissive to *S*. Typhi growth (Figure 3A). The analysis of HPS-1 knockdown iBMDMs infected with *S*. Typhi by flow cytometry also revealed that in HPS-1 depleted cells there was a higher percentage of cells that exhibited high red fluorescence (5.7%), i.e. contained more intracellular bacteria, compared to control cells (1.15%) (Figure 3B). Overall, the results obtained showed that iBMDMs could be used as a suitable cellular model to validate the candidates from the shRNA screen.

**Figure 3.**
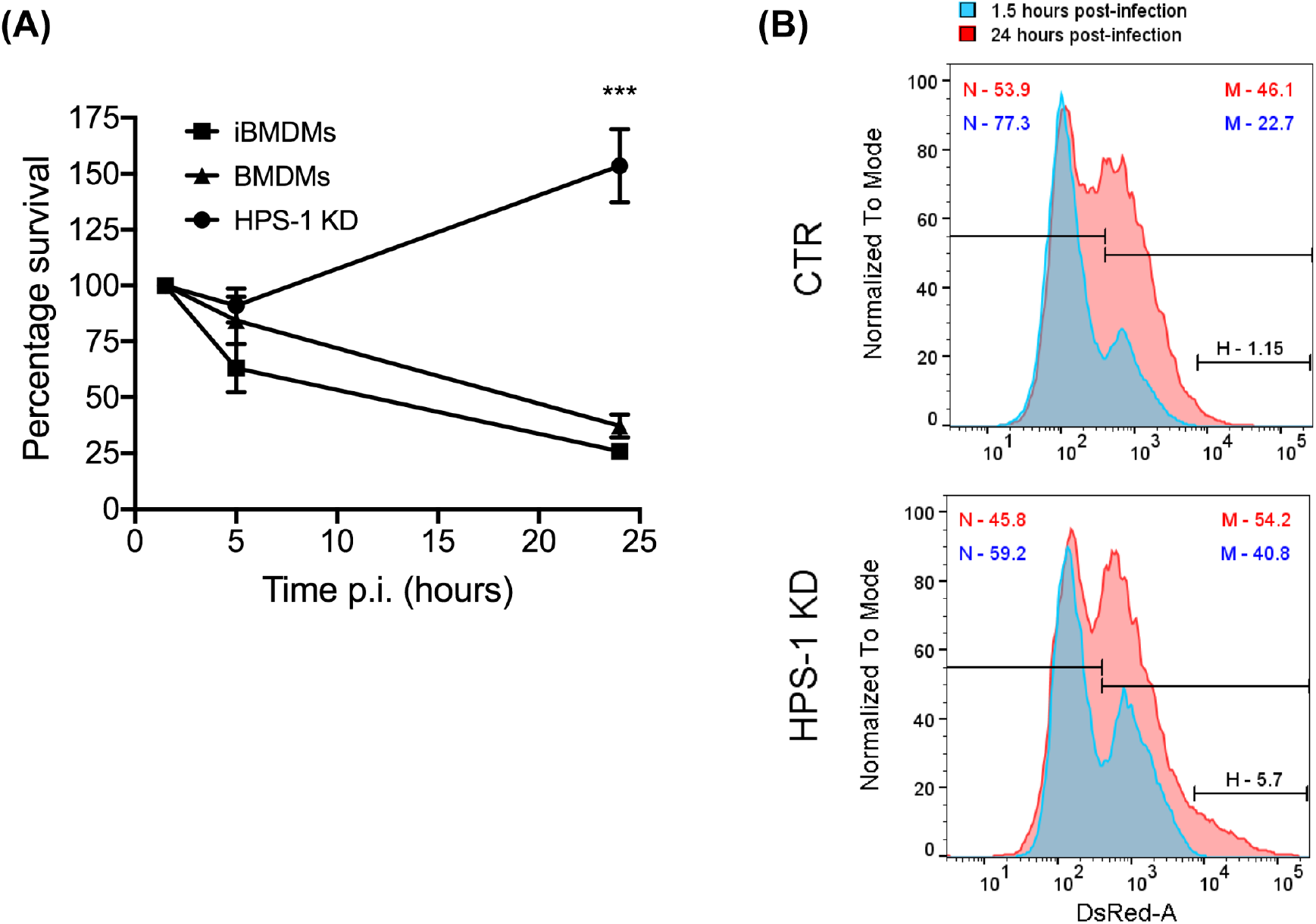
Depletion of HPS-1 allows *S*. Typhi to survive better in immortalised bone-marrow-derived macrophages (iBMDMs). (A) BMDMs, iBMDMs or iBMDMs HPS-1 knockdown (HPS-1 KD) were infected with S. Typhi wild-type (moi 5). At the indicated times post-infection, cells were lysed and CFU enumerated. The CFUs obtained are normalised against the first time point 1.5 hpi, which is considered as 100% of uptake. Values are means ± SEM of two independent experiments performed in triplicates. *P* values were determined by the Student’s *t* test: *** *P*<0.005. (B) iBMDMs transduced with lentiviruses encoding either non-coding sequences (CTR) or an shRNA targeting HPS-1 (HPS-1 KD) were infected with *S*. Typhi::mCherry. At different times post-infection (1.5 or 24 hpi) cells were fixed and analysed by flow cytometry. The abundance in percentage of the different subpopulations is indicated.

For each candidate gene we took the second best shRNA (see Table S4) and we checked the levels of knockdown achieved in iBMDM (Figure S5). The reduction in mRNA expression was over 50% in all cases except for Rab27 and Rab40c. In addition, Rab25 mRNA was not detectable. This suggested that these three Rabs were false positives and were removed from our validation list. We then measured the CFU 24 h after infection with *S*. Typhi::*mCherry* (Figure 4). As expected, we confirmed that depletion of Rab32 and HPS-1 in mouse macrophages lead to an increased survival of *S*. Typhi (Spano and Galan., 2012). Among the other 7 Rab GTPases tested, only the knockdown of Rab1b showed a significant increase of *S*. Typhi survival (Figure 4). To further confirm that the phenotype observed with the shRNA targeting Rab1b#2 (z-score = 1.69) was not due to off-target effect of that particular shRNA, we generated lentivirus particles encoding the best scored Rab1b shRNA (Rab1b#1; z-score=1.78). The results obtained were similar to those reported above and are shown in Figure 4.

**Figure 4.**
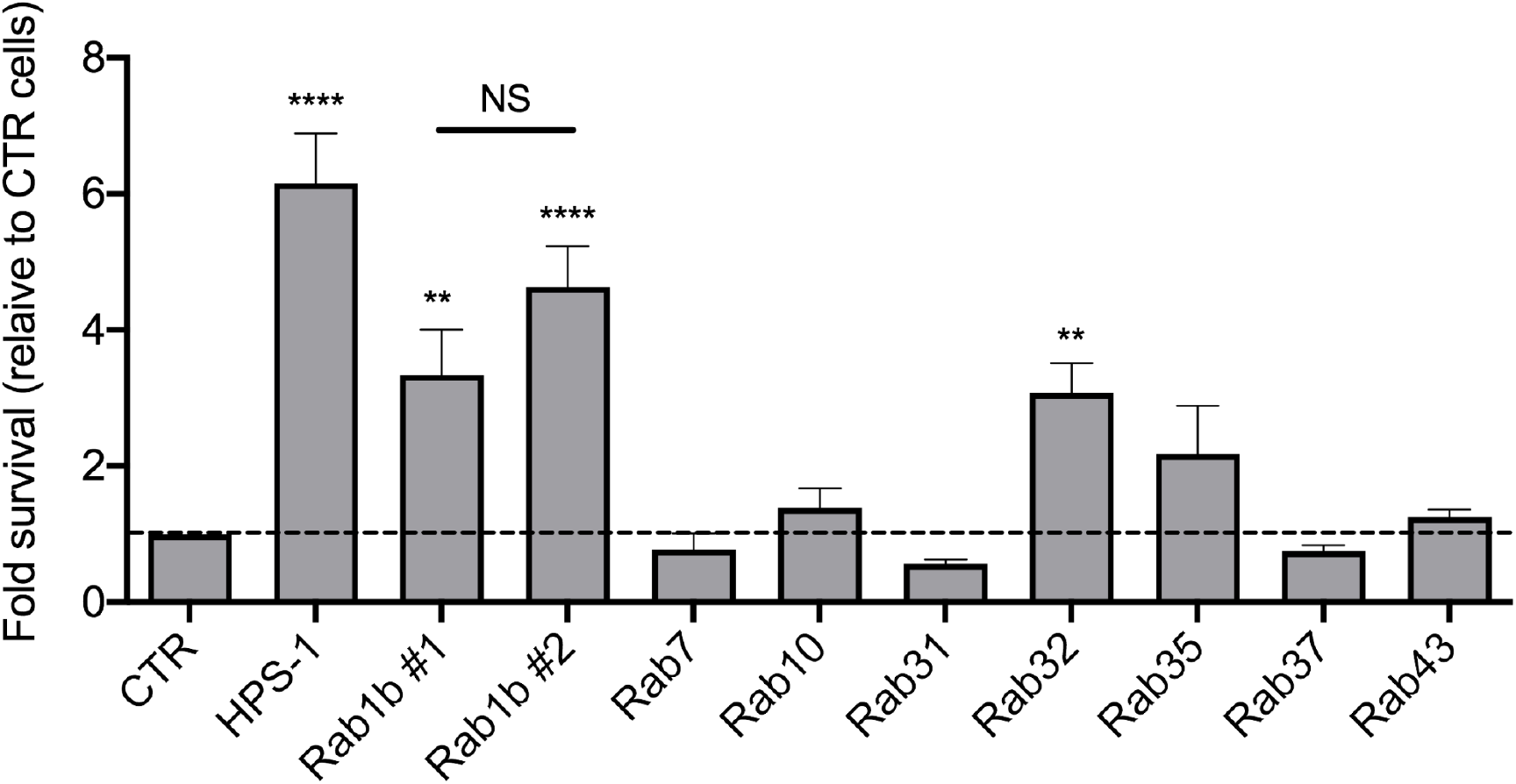
Validation of candidate genes. For each gene, lentivirus particles were generated and used to transduced iBMDMs. After selection with 5 μg/ml puromycin for 48 hours, cells were infected with *S*. Typhi. At different times post-infection (1.5 and 24 hours), cells were lysed and CFUs enumerated. The CFUs obtained were normalized against 1.5 hpi and the data represent the fold-survival relative to CTR cells. Values are means ± SEM of at least two independent experiments performed in triplicates. *P* values were determined by the Student’s t test: \**P<0.05*; \*\**P<0.01*; \*\*\**P<0.005*; \*\*\*\**P<0,001*.

## Discussion

Over the past years, unbiased genetic screenings aimed to understand the interaction of bacterial pathogens with their eukaryotic hosts have been crucial to identify host genes that play important roles in dealing with infections (Zhou et al., 2012, Yeung et al., 2019, Li et al., 2017). However, as primary macrophages undergo terminal differentiation and cannot be propagated in culture, the majority of these screens have been performed in alternative cell lines. Immortalized cell lines have the advantages of being easier to manipulate genetically and more straightforward to expand in culture, but their continuous subculture can lead to altered macrophage functions, including pathogen killing, limiting their suitability for screening experiments.

In this study, we performed a loss-of-function screen in primary mouse macrophages using viral delivered shRNAs in the context of *S*. Typhi infection. To the best of our knowledge, this screen is the first of its kind. We have shown that shRNAs, delivered using lentiviral particles, represent a good method to achieve sufficient level of knockdown for further phenotypic analyses. Most importantly, we have developed a protocol that allows sorting fixed cells and extracting good quality DNA from fixed cells for further next-generation sequencing. This is important due to the restrictions of working with *S*. Typhi, which in many places requires containment level 3 (CL3) facilities. Therefore by fixing the samples and sorting them out of the CL3 we eliminated the requirement of a cell sorter in CL3 environment.

Our fluorescence microscopy and flow cytometry analyses of HPS-1-depleted macrophages (our positive control) in response to *S*. Typhi infection showed that the removal of HPS-1 led to a population of macrophages containing a very high number of bacteria (>6), but also induced a shift in the intermediate population with more cells containing between 3 and 6 bacteria (Figure 2C and S2). Therefore in our sorting strategy we selected a gate including both the very low abundant population containing very high number and the upper limit of the population containing lower numbers of intracellular bacteria (gate H in Figure 4A). This slightly more inclusive gating also allowed us to reduce the total number of cells that needed to be sorted in order to recover enough cells from the H population for further analyses. The down side of this expansion is that we have increased the noise and probably the number of false positives. These aspects should be taken into consideration when planning for screens with larger libraries. Alternatives, such as performing a higher stringency secondary screen with customized libraries targeting the candidate genes identified with a first low stringent strategy, should be carefully considered.

In addition to providing a proof-of-concept study, showing that viral delivery of small shRNA in primary macrophage is effective and results in a measurable phenotype, our screen revealed that Rab1b is important to control the growth of *S*. Typhi in mouse macrophages. Rab1 is known to regulate vesicle transport from the endoplasmic reticulum (ER) to the Golgi complex (Plutner et al., 1991). In addition, Rab1b is involved in autophagy, a process by which the cell delivers cytosolic material (misfolded proteins, damaged organelles and pathogens) to the lysosome for degradation (Kakuta et al., 2017). In the context of bacterial infections, intracellular pathogens such as *Legionella pneumophila* and *Yersinia pestis* actively recruit Rab1 to the pathogen-containing vacuoles to prevent acidification of this compartment allowing the bacteria to survive inside macrophages (Kagan et al., 2004, Connor et al., 2015). In other cases, Rab1 is blocked to prevent autophagy and therefore, pathogen killing. This is the case of *S*. Typhimurium, which uses two effectors, SseF and SseG to block Rab1 to prevent autophagy and bacterial clearance (Feng et al., 2018). Two more recent studies have identified that *S*. Typhimurium uses a third effector, SseK3, to covalently modify Rab1. The result of this modification impairs ER-Golgi trafficking limiting the secretion of cytokines during *Salmonella* infection (Gan et al., 2020, Meng et al., 2020). The fact that a single pathogen uses multiple effectors to target a single host protein highlights the importance of Rab1 in the context of *Salmonella* infection. SseK3 is absent from *S*. Typhi and while SseG is highly conserved, SseF has evolved differently in different *Salmonella* serovars (Eswarappa et al., 2008). These differences in the arsenal of effectors between these two serovars could explain why *S*. Typhi cannot block Rab1b in mouse cells.

In summary, we have described experimental conditions to perform loss-of-function screens in primary macrophages negotiating the constraints required by a CL3 environment when working with certain pathogens. This offers the potential to perform unbiased large-scale screenings in primary macrophages to identify novel mechanisms of pathogen killing. Interestingly, as a result of a targeted, small-scale screen we have identified a role of Rab1b in controlling the growth of the human pathogen *S*. Typhi in mouse macrophages. To the best of our knowledge, this is the first report showing a Rab GTPase other than Rab32 having a role in the restriction of *S*. Typhi in mice. Future work will define how Rab1b prevents *S*. Typhi growth in mouse macrophages and whether this restriction mechanism is related to other previously described macrophage-killing mechanisms against this human pathogen.

**This paper is part of Stefania Spanò’s scientific legacy and this would have not been possible without her intelligence, vision and persistence. A dreadful destiny has snatched her from us too early, but her discoveries and ideas are living and flourishing.**

## Author contribution

VSC performed the experiments and wrote the manuscript with input from MB and DC. RC performed experiments. MB, VSC and SS contributed to the design of the study and the analysis of the results. All authors contributed to the final version of the manuscript.

## Acknowledgements

We are very grateful to Leigh Knodler for her generous gift of P22 phages from a *S*. Typhimurium *glmS::Cm::mCherry* strain. We thank member of the Spanò/Baldassarre laboratory for their feedback throughout this project.

## References

1. Weiss G, Schaible UE. Macrophage defense mechanisms against intracellular bacteria. Immunol Rev (2015) 264:182–203 doi: 10.1111/imr.12266 [doi].

2. Fields PI, Swanson RV, Haidaris CG, Heffron F. Mutants of Salmonella typhimurium that cannot survive within the macrophage are avirulent. Proc Natl Acad Sci U S A (1986) 83:5189–93 doi: 10.1073/pnas.83.14.5189 [doi].

3. Jennings E, Thurston TLM, Holden DW. Salmonella SPI-2 Type III Secretion System Effectors: Molecular Mechanisms And Physiological Consequences. Cell Host Microbe (2017) 22:217–31 doi: S1931-3128(17)30292-5 [pii].

4. Galan JE, Lara-Tejero M, Marlovits TC, Wagner S. Bacterial type III secretion systems: specialized nanomachines for protein delivery into target cells. Annu Rev Microbiol (2014) 68:415–38 doi: 10.1146/annurev-micro-092412-155725 [doi].

5. Spano S, Gao X, Hannemann S, Lara-Tejero M, Galan JE. A Bacterial Pathogen Targets a Host Rab-Family GTPase Defense Pathway with a GAP. Cell Host Microbe (2016) 19:216–26 doi: 10.1016/j.chom.2016.01.004 [doi].

6. Dougan G, Baker S. Salmonella enterica serovar Typhi and the pathogenesis of typhoid fever. Annu Rev Microbiol (2014) 68:317–36 doi: 10.1146/annurev-micro-091313-103739 [doi].

7. Gibani MM, Britto C, Pollard AJ. Typhoid and paratyphoid fever: a call to action. Curr Opin Infect Dis (2018) 31:440–8 doi: 10.1097/QCO.0000000000000479 [doi].

8. Spano S, Galan JE. A Rab32-dependent pathway contributes to Salmonella typhi host restriction. Science (2012) 338:960–3 doi: 10.1126/science.1229224 [doi].

9. Stein MP, Muller MP, Wandinger-Ness A. Bacterial pathogens commandeer Rab GTPases to establish intracellular niches. Traffic (2012) 13:1565–88 doi: 10.1111/tra.12000 [doi].

10. McGourty K, Thurston TL, Matthews SA, Pinaud L, Mota LJ, Holden DW. Salmonella inhibits retrograde trafficking of mannose-6-phosphate receptors and lysosome function. Science (2012) 338:963–7 doi: 10.1126/science.1227037 [doi].

11. Bakowski MA, Braun V, Lam GY, Yeung T, Heo WD, Meyer T, et al. The phosphoinositide phosphatase SopB manipulates membrane surface charge and trafficking of the Salmonella-containing vacuole. Cell Host Microbe (2010) 7:453–62 doi: 10.1016/j.chom.2010.05.011 [doi].

12. Spano S, Galan JE. Taking control: Hijacking of Rab GTPases by intracellular bacterial pathogens. Small GTPases (2018) 9:182–91 doi: 10.1080/21541248.2017.1336192 [doi].

13. Hardiman CA, McDonough JA, Newton HJ, Roy CR. The role of Rab GTPases in the transport of vacuoles containing Legionella pneumophila and Coxiella burnetii. Biochem Soc Trans (2012) 40:1353–9 doi: 10.1042/BST20120167 [doi].

14. Seto S, Tsujimura K, Koide Y. Rab GTPases regulating phagosome maturation are differentially recruited to mycobacterial phagosomes. Traffic (2011) 12:407–20 doi: 10.1111/j.1600-0854.2011.01165.x [doi].

15. Galan JE, Curtiss R,3rd. Distribution of the invA, -B, -C, and -D genes of Salmonella typhimurium among other Salmonella serovars: invA mutants of Salmonella typhi are deficient for entry into mammalian cells. Infect Immun (1991) 59:2901–8.

16. Baldassarre M, Solano-Collado V, Balci A, Colamarino R, Dambuza I, Reid D, et al. The Rab32/BLOC-3 dependent pathway mediates host-defence against different pathogens in human macrophages. Science Advances (2020) doi: in press.

17. Brennan MA, Cookson BT. Salmonella induces macrophage death by caspase-1-dependent necrosis. Mol Microbiol (2000) 38:31–40 doi: mmi2103 [pii].

18. Weischenfeldt J, Porse B. Bone Marrow-Derived Macrophages (BMM): Isolation and Applications. CSH Protoc (2008) 2008:pdb.prot5080 doi: 10.1101/pdb.prot5080 [doi].

19. Shi X, Mihaylova VT, Kuruvilla L, Chen F, Viviano S, Baldassarre M, et al. Loss of TRIM33 causes resistance to BET bromodomain inhibitors through MYC- and TGF-beta-dependent mechanisms. Proc Natl Acad Sci U S A (2016) 113:E4558–66 doi: 10.1073/pnas.1608319113 [doi].

20. Carette JE, Guimaraes CP, Wuethrich I, Blomen VA, Varadarajan M, Sun C, et al. Global gene disruption in human cells to assign genes to phenotypes by deep sequencing. Nat Biotechnol (2011) 29:542–6 doi: 10.1038/nbt.1857 [doi].

21. Silva JM, Marran K, Parker JS, Silva J, Golding M, Schlabach MR, et al. Profiling essential genes in human mammary cells by multiplex RNAi screening. Science (2008) 319:617–20 doi: 10.1126/science.1149185 [doi].

22. Thompson CD, Frazier-Jessen MR, Rawat R, Nordan RP, Brown RT. Evaluation of methods for transient transfection of a murine macrophage cell line, RAW 264.7. BioTechniques (1999) 27:824,6, 828–30, 832 doi: 10.2144/99274rr05 [doi].

23. Guiet R, Verollet C, Lamsoul I, Cougoule C, Poincloux R, Labrousse A, et al. Macrophage mesenchymal migration requires podosome stabilization by filamin A. J Biol Chem (2012) 287:13051–62 doi: 10.1074/jbc.M111.307124 [doi].

24. Naldini L, Blomer U, Gallay P, Ory D, Mulligan R, Gage FH, et al. In vivo gene delivery and stable transduction of nondividing cells by a lentiviral vector. Science (1996) 272:263–7 doi: 10.1126/science.272.5259.263 [doi].

25. Schroers R, Sinha I, Segall H, Schmidt-Wolf IG, Rooney CM, Brenner MK, et al. Transduction of human PBMC-derived dendritic cells and macrophages by an HIV-1-based lentiviral vector system. Mol Ther (2000) 1:171–9 doi: 10.1006/mthe.2000.0027 [doi].

26. Zeng L, Yang S, Wu C, Ye L, Lu Y. Effective transduction of primary mouse blood- and bone marrow-derived monocytes/macrophages by HIV-based defective lentiviral vectors. (2006) doi: doi:10.1016/j.jviromet.2005.12.006.

27. Miller MR, Blystone SD. Reliable and inexpensive expression of large, tagged, exogenous proteins in murine bone marrow-derived macrophages using a second generation lentiviral system. J Biol Methods (2015) 2:e23 doi: 10.14440/jbm.2015.66 [doi].

28. Zhou H, DeLoid G, Browning E, Gregory DJ, Tan F, Bedugnis AS, et al. Genome-wide RNAi screen in IFN-gamma-treated human macrophages identifies genes mediating resistance to the intracellular pathogen Francisella tularensis. PLoS One (2012) 7:e31752 doi: 10.1371/journal.pone.0031752 [doi].

29. Yeung ATY, Choi YH, Lee AHY, Hale C, Ponstingl H, Pickard D, et al. A Genome-Wide Knockout Screen in Human Macrophages Identified Host Factors Modulating Salmonella Infection. mBio (2019) 10:10.1128/mBio.02169–19 doi: e02169-19 [pii].

30. Li N, Katz S, Dutta B, Benet ZL, Sun J, Fraser ID. Genome-wide siRNA screen of genes regulating the LPS-induced NF-kappaB and TNF-alpha responses in mouse macrophages. Sci Data (2017) 4:170008 doi: 10.1038/sdata.2017.8 [doi].

31. Plutner H, Cox AD, Pind S, Khosravi-Far R, Bourne JR, Schwaninger R, et al. Rab1b regulates vesicular transport between the endoplasmic reticulum and successive Golgi compartments. J Cell Biol (1991) 115:31–43 doi: 10.1083/jcb.115.1.31 [doi].

32. Kakuta S, Yamaguchi J, Suzuki C, Sasaki M, Kazuno S, Uchiyama Y. Small GTPase Rab1B is associated with ATG9A vesicles and regulates autophagosome formation. FASEB J (2017) 31:3757–73 doi: 10.1096/fj.201601052R [doi].

33. Kagan JC, Stein MP, Pypaert M, Roy CR. Legionella subvert the functions of Rab1 and Sec22b to create a replicative organelle. J Exp Med (2004) 199:1201–11 doi: 10.1084/jem.20031706 [doi].

34. Connor MG, Pulsifer AR, Price CT, Abu Kwaik Y, Lawrenz MB. Yersinia pestis Requires Host Rab1b for Survival in Macrophages. PLoS Pathog (2015) 11:e1005241 doi: 10.1371/journal.ppat.1005241 [doi].

35. Feng ZZ, Jiang AJ, Mao AW, Feng Y, Wang W, Li J, et al. The Salmonella effectors SseF and SseG inhibit Rab1A-mediated autophagy to facilitate intracellular bacterial survival and replication. J Biol Chem (2018) 293:9662–73 doi: 10.1074/jbc.M117.811737 [doi].

36. Gan J, Scott NE, Newson JPM, Wibawa RR, Wong Fok Lung T, Pollock GL, et al. The Salmonella Effector SseK3 Targets Small Rab GTPases. Front Cell Infect Microbiol (2020) 10:419 doi: 10.3389/fcimb.2020.00419 [doi].

37. Meng K, Zhuang X, Peng T, Hu S, Yang J, Wang Z, et al. Arginine GlcNAcylation of Rab small GTPases by the pathogen Salmonella Typhimurium. Commun Biol (2020) 3:287,020–1005-2 doi: 10.1038/s42003-020-1005-2 [doi].

38. Eswarappa SM, Janice J, Nagarajan AG, Balasundaram SV, Karnam G, Dixit NM, et al. Differentially evolved genes of Salmonella pathogenicity islands: insights into the mechanism of host specificity in Salmonella. PLoS One (2008) 3:e3829 doi: 10.1371/journal.pone.0003829 [doi].

