## Supplementary Figures and Tables S1, S4 and S5 for "A small-scale shRNA screen in primary mouse macrophages identifies a role for the Rab GTPase Rab1b in controlling *Salmonella* Typhi growth"

This file contains Supplementary Figures S1-S4 and Supplementary Tables S1, S4, S5

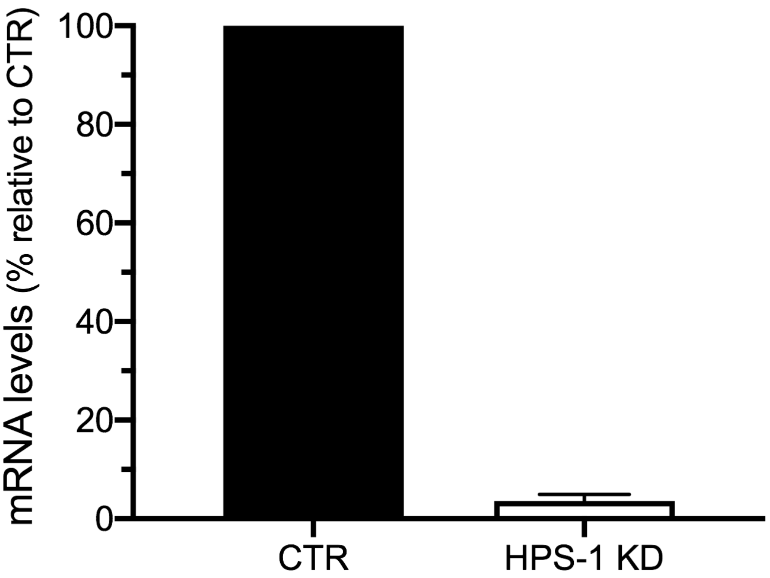


**Figure S1.** mRNA levels of HPS-1 in BMDMs. BMDMs were transduced with an shRNA targeting HPS-1 (HPS-1 KD) or non-targeting sequences (CTR) and the transcript levels of HPS-1 were determined by RT-qPCR. The GAPDH gene was used as reference.

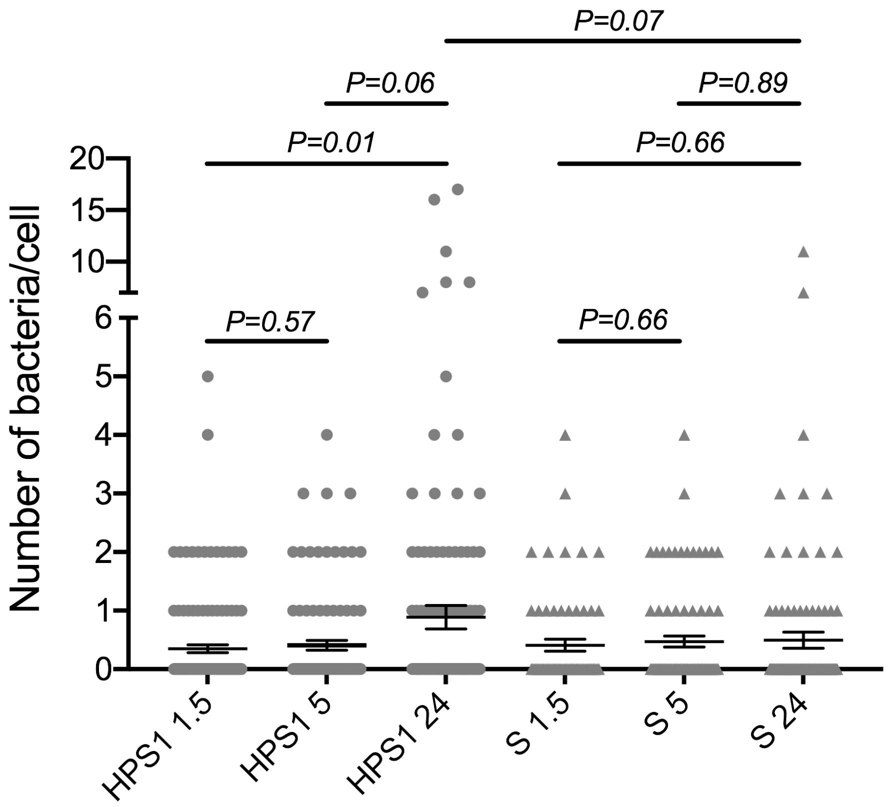


**Figure S2**. BMDMs HPS-1 Knockdown (HPS1) or control (Scrambled shRNA; S) were infected with *S*. Typhi::*mCherry* (moi 10). At different times post-infection, cells were fixed with 4% PFA and analysed by fluorescence microscopy to determine the number of bacteria per cell. Student´s t-test (unpaired, two tailed) was used to calculate statistical significance of the data and the *P* values obtained are indicated.

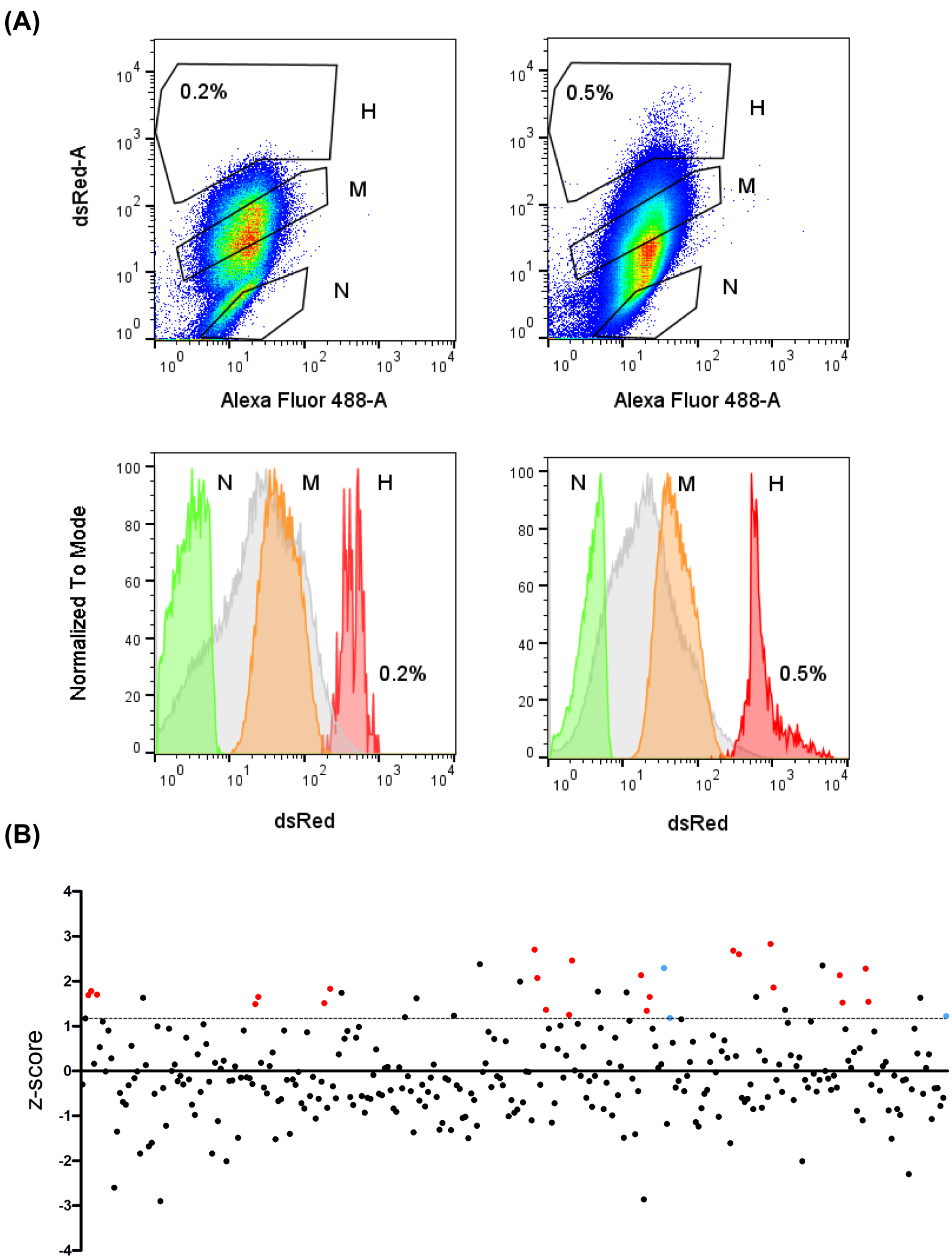


**Figure S3**. Cell sorting by flow cytometry and analysis of the results (z-score) **(A)** Samples ar 1.5 and 24 hours-post infection were analyzed and sorted in a BD Influx BSLII Sorter (Ian Fraser Cytometry Centre, University of Aberdeen). For a more accurate separation of the different cellular subpopulations, cell auto-fluorescence (Alexa Fluor 488-A) was used together with bacteria fluorescence (DsRed) to select sorting gates. Sorting gates used to isolate the different cell sub-populations are indicated as well as the percentage of cells carrying high amount of intracellular bacteria. The subpopulations sorted are shown based in DsRed fluorescence in the lower panel. N (non-infected), M (infected cells with low number of intracellular bacteria), H (infected cells with high number of intracellular bacteria) **(B)** Dot-plot of z-scores. Genes exhibiting more than 1 hairpin with a z-score >1.18 are represented in red. Rab32 and HPS-1 (positive controls) are represented in blue.

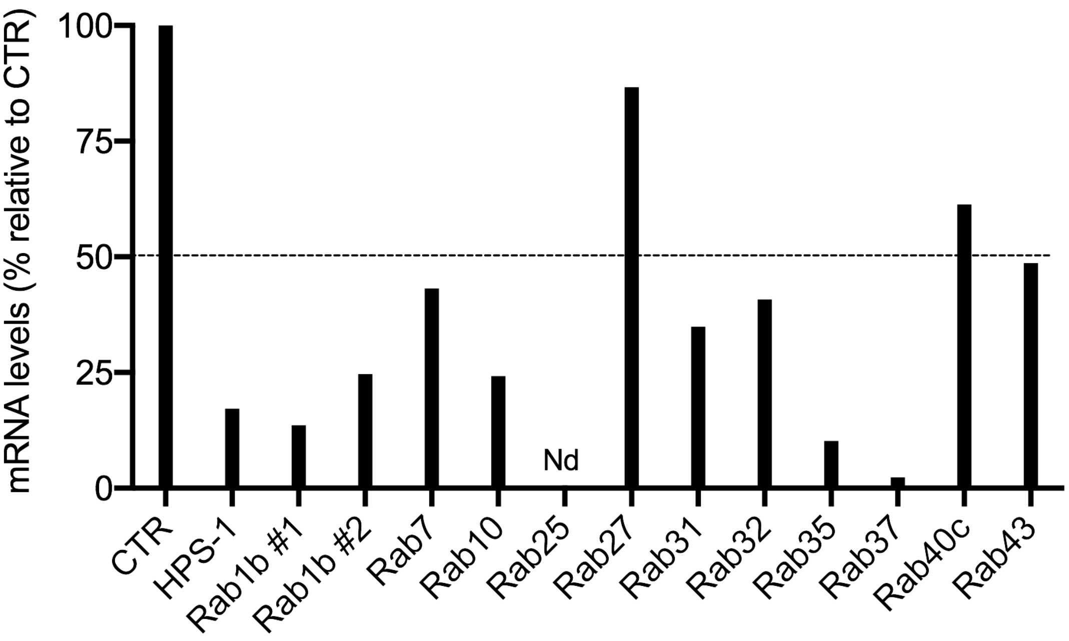


**Figure S4.** mRNA levels of Rab GTPases validated. iBMDMs were transduced with the second best shRNA of each gene or non-targeting sequences (CTR) and the transcript levels of each gene were determined by RT-qPCR. The GAPDH gene was used as reference.

**Supplementary Tables**

**Table S1.** shRNA library comprising the mouse Rab GTPases and HPS-1 (positive control)

| GENE | TRCN_ID | shRNA SEQUENCE (SENSE STRAND) |
|---|---|---|
| Rab1b | TRCN0000100822 | GATTTCAAGATTCGAACCATT |
| Rab1b | TRCN0000100823 | CATGGCATCATTGTGGTGTAT |
| Rab1b | TRCN0000302710 | CCACGCACCTTTCTTTGGAAT |
| Rab1b | TRCN0000302711 | GCCAAGAATGCCACCAATGTT |
| Rab1b | TRCN0000302786 | GCGGTTTGCTGATGACACTTA |
| Rab1b | TRCN0000379948 | ACACCACGGCCAAGGAATTTG |
| Rab1b | TRCN0000381762 | AGTCCTACGCCAACGTGAAAC |
| Rab2b | TRCN0000100406 | ACGGCCTTATATTCATGGAAA |
| Rab2b | TRCN0000100408 | AGGTTTATTTGATGTCCACAA |
| Rab2b | TRCN0000287555 | GCAGTCAATTACCTCCTCAGT |
| Rab2b | TRCN0000287556 | CCTCCTTCAGTTTACCGACAA |
| Rab2b | TRCN0000287557 | CCGCGTAGATTATGGCTCTTT |
| Rab3a | TRCN0000089148 | GCCTTATACTTTGGGATAAAT |
| Rab3a | TRCN0000089149 | CGACTATATGTTCAAGATCCT |
| Rab3a | TRCN0000089150 | ACCAATGAGGAGTCATTTAAT |
| Rab3a | TRCN0000089151 | TCACCAATGAGGAGTCATTTA |
| Rab3a | TRCN0000089152 | CAGCGCCAAGGACAACATTAA |
| Rab3b | TRCN0000089363 | GCAGTCTTTGATAAACTGTAA |
| Rab3b | TRCN0000089364 | GCAGCAGAACTGCTCTTGTTA |
| Rab3b | TRCN0000089366 | CGATAAGATGTCTGACTCGAT |
| Rab3b | TRCN0000089367 | CTTCAAAGTGAAGACAGTCTA |
| Rab3c | TRCN000008945 | CCTGTGTTAATATGTGGCAAA |

| | | |
|---|---|---|
| | 3 | |
| Rab3c | TRCN0000089454 | CGATTCCTTTACATCTGCATT |
| Rab3c | TRCN0000089455 | GATAACATCAACGTGAAGCAA |
| Rab3c | TRCN0000089456 | CCCAGGTTATCCTGGCTGGAA |
| Rab3c | TRCN0000089457 | TGTTCCGTTATGCCGACGATT |
| Rab3d | TRCN0000089443 | CGGTGCAAATTATGTTCTTAA |
| Rab3d | TRCN0000089444 | CAAACTGCTCTTGATCGGGAA |
| Rab3d | TRCN0000089445 | CATCATCTGTGACAAGATGAA |
| Rab3d | TRCN0000089446 | CTACCGACATGACAAGAGGAT |
| Rab3d | TRCN0000089447 | CCGACATGACAAGAGGATCAA |
| Rab4a | TRCN0000088973 | GCTTTCTCATTGCGTTGGTTA |
| Rab4a | TRCN0000088974 | GATAATAAATGTCGGTGGTAA |
| Rab4a | TRCN0000088975 | AGATGACTCAAATCATACCAT |
| Rab4a | TRCN0000088976 | CCTACAATGCGCTTACTAATT |
| Rab4a | TRCN0000088977 | CTCAAATCATACCATAGGAAT |
| Rab4b | TRCN0000089433 | CCACCAGTTTATTGAGAATAA |
| Rab4b | TRCN0000089434 | GCATTCAGTATGGCGACATAT |
| Rab4b | TRCN0000089435 | TCCACCAGTTTATTGAGAATA |
| Rab4b | TRCN0000089436 | AGACTGTGAAACTACAGATTT |
| Rab4b | TRCN0000089437 | CATACAACTCACTCGCTGCTT |
| Rab5a | TRCN0000100795 | CCCAAGCAAATGGTGTAATTT |
| Rab5a | TRCN0000100796 | GCAGCCATAGTTGTGTATGAT |
| Rab5a | TRCN0000100797 | CGCTTTGTGAAAGGCCAATTT |
| Rab5a | TRCN0000100798 | GCTGGTCAAGAACGGTATCAT |
| Rab5a | TRCN0000100799 | CAAGCAGCCATAGTTGTGTAT |

| | | |
|---|---|---|
| Rab5b | TRCN0000100620 | GCACTTTAATTGATGGTAGTT |
| Rab5b | TRCN0000100621 | CCGTGTGTTTAGATGACACAA |
| Rab5b | TRCN0000100622 | CCTGGCAATAGCAAAGAAGTT |
| Rab5b | TRCN0000100623 | GCAGATGACAACAGCTTATTA |
| Rab5b | TRCN0000100624 | CGTGGTCTATGATATTACTAA |
| Rab5c | TRCN0000100745 | CCCGACTGGAATCTACTCTAA |
| Rab5c | TRCN0000100746 | CCCAACATCGTCATTGCACTA |
| Rab5c | TRCN0000100747 | GCAATGAACGTGAATGAAATT |
| Rab5c | TRCN0000100748 | GCAACAAGATCTGTCAGTTTA |
| Rab5c | TRCN0000100749 | GCTAAGAAGCTTCCCAAGAAT |
| Rab6b | TRCN0000100900 | CCACAGTCAAATCCAACTTTA |
| Rab6b | TRCN0000100901 | CGGGATTGACTTCTTGTCAAA |
| Rab6b | TRCN0000100902 | CCAGCAGACTTCTAAATGGAT |
| Rab6b | TRCN0000100903 | CTGTGGTGGTATATGACATTA |
| Rab6b | TRCN0000100904 | GCTGTGGTGGTATATGACATT |
| Rab7 | TRCN0000100880 | GCGGCAGTATTCTGTACAGTA |
| Rab7 | TRCN0000100881 | TGAACCCATCAAACTGGACAA |
| Rab7 | TRCN0000100882 | GAAGTTCAGTAACCAGTACAA |
| Rab7 | TRCN0000100883 | TGCTGTGTTCTGGTGTTTGAT |
| Rab7 | TRCN0000100884 | GCCCTTAAACAGGAAACAGAA |
| Rab8a | TRCN0000100420 | CCATGAAATGAATCTGTCTTT |
| Rab8a | TRCN0000100421 | CGCCTTCAACTCCACATTCAT |
| Rab8a | TRCN0000100422 | CGGAATTGGATTCGGAACATT |
| Rab8a | TRCN0000100423 | CTACGACATTACCAATGAGAA |
| Rab8a | TRCN000010042 | CTCGATGGCAAGAGGATTAAA |

| | | |
|---|---|---|
| | 4 | |
| Rab8b | TRCN000010053<br>5 | GCCAAGAACTAACAGAACTTT |
| Rab8b | TRCN000010053<br>6 | CCGAACAATTACGACAGCATA |
| Rab8b | TRCN000010053<br>7 | CCGGTCTAAGAAGACCAGTTT |
| Rab8b | TRCN000010053<br>8 | CAGGAAAGATTCCGAACAATT |
| Rab8b | TRCN000010053<br>9 | CAAATGTGATATGAACGACAA |
| Rab8b | TRCN000029536<br>3 | AGTGCAAAGTCGAGTACAAAT |
| Rab8b | TRCN000029542<br>4 | TTTACACTTGCACGGGATATA |
| Rab8b | TRCN000029542<br>5 | AGAAGTTAGCAATTGACTATG |
| Rab8b | TRCN000038034<br>3 | ATAACAGAGAGCCGGTCTAAG |
| Rab9b | TRCN000010066<br>0 | GCCAAGTAGTTTATGTCATTT |
| Rab9b | TRCN000010066<br>1 | CCGTTATGTAACCAACAAGTT |
| Rab9b | TRCN000010066<br>2 | GCAGATGTAAAGGACCCAGAT |
| Rab9b | TRCN000010066<br>3 | GAGCCTTAGAACACCATTCTA |
| Rab9b | TRCN000010066<br>4 | GTAGAGGAACAGCTGGAACAT |
| Rab10 | TRCN000010083<br>5 | GCCTATTAACTGTCAGTTAAT |
| Rab10 | TRCN000010083<br>9 | CATGCCAATGAAGATGTGGAA |
| Rab10 | TRCN000033554<br>3 | CGATGCCTTCAATACCACCTT |
| Rab10 | TRCN000033554<br>4 | GAGAGTTGTACCGAAAGGCAA |
| Rab10 | TRCN000033562<br>3 | GCGTTCCTCACATTAGCTGAA |
| Rab10 | TRCN000034857<br>3 | GACTGCTTGCGGACTATTATA |
| Rab11a | TRCN000010034<br>0 | CCCTGTAAACATAACAGCATT |
| Rab11a | TRCN000010034<br>2 | CAGGGCTATAACGTCTGCATA |
| Rab11a | TRCN000010034<br>3 | ACCTCTTTAAAGTTGTCCTTA |
| Rab11a | TRCN000010034<br>4 | CAGAGATATACCGCATTGTTT |

| | | |
|---|---|---|
| Rab11a | TRCN0000305795 | AGTAGGTGCCTTATTGGTTTA |
| Rab11a | TRCN0000305864 | TAACCTCCTGTCTCGATTTAC |
| Rab11a | TRCN0000324936 | GAGAGATCATGCTGATAGTAA |
| Rab11b | TRCN0000100255 | CCTGTCTGCAAGTGAAGCAAT |
| Rab11b | TRCN0000100256 | CCTCACAGAAATCTACCGTAT |
| Rab11b | TRCN0000100257 | CCTATTCAAAGTGGTGCTTAT |
| Rab11b | TRCN0000100258 | CAAGACCATCAAGGCTCAGAT |
| Rab11b | TRCN0000100259 | CCTAGAGAGCAAGAGTACCAT |
| Rab13 | TRCN0000100855 | GAGCATTTCTTGCCTCCTATT |
| Rab13 | TRCN0000100856 | GCCAAGAACGATTCAAGACAA |
| Rab13 | TRCN0000100857 | CCAAGAACGATTCAAGACAAT |
| Rab13 | TRCN0000100858 | CGAGAGCACAGAATCCGATTT |
| Rab13 | TRCN0000100859 | GATGAGAAATCCTTCGAGAAT |
| Rab14 | TRCN0000089374 | GCACAGAGAGATGTTACCTAT |
| Rab14 | TRCN0000089376 | CCAATCCAAACACTGTAATAA |
| Rab14 | TRCN0000306372 | ATCACCAGAAGAAGTACATAT |
| Rab14 | TRCN0000306430 | CGGTTACACGGAGCTACTATA |
| Rab14 | TRCN0000332297 | ACCAATCCAAACACTGTAATA |
| Rab14 | TRCN0000332372 | CAGAGAGATGTTACCTATGAA |
| Rab14 | TRCN0000332373 | CCTTTGATTGTCCTTGTGATA |
| Rab15 | TRCN0000093169 | CCCTGAAGTATAGCAACAGAA |
| Rab15 | TRCN0000093170 | CCAGACTATCACAAAGCAGTA |
| Rab15 | TRCN0000093171 | GCATGGACTTCTACGAAACAA |
| Rab15 | TRCN0000093172 | CGGTGTTGACTTTAAGATGAA |
| Rab15 | TRCN000009317 | GCGCTCCTATCAGCATATCAT |

| | | |
|---|---|---|
| | 3 | |
| Rab17 | TRCN000010091 5 | GCTGCCTCTTTGTCCATTCAT |
| Rab17 | TRCN000010091 6 | CCGGTACATGAAGCAGGACTT |
| Rab17 | TRCN000010091 7 | TCTGAGATCTTCAACACTGTT |
| Rab17 | TRCN000010091 8 | CTCCTGGTTTATGACATCACT |
| Rab17 | TRCN000010091 9 | GCGCCAGTGCTGTGCACGATA |
| Rab18 | TRCN000017702 8 | GATGGAAATAAGGCTAAACTT |
| Rab18 | TRCN000017779 4 | CGATAGAAATGAAGGCTTGAA |
| Rab18 | TRCN000017793 7 | GCAACAATAGGTGTTGACTTT |
| Rab18 | TRCN000017829 1 | GCTAAACTTGCAATATGGGAT |
| Rab18 | TRCN000018181 2 | GAACCAGAACAAAGGAGTCAA |
| Rab18 | TRCN000018211 1 | CCCAGCTATTATAGAGGTGCA |
| Rab18 | TRCN000019749 9 | CAGGGAGTTATATTAGTCTAT |
| Rab18 | TRCN000021533 1 | CAAGAGAGGTTCAGAACATTA |
| Rab18 | TRCN000021587 9 | CTTGTTGAGAAGATCATTCAG |
| Rab18 | TRCN000021662 2 | GTAAACATGCTAGTTGGAAAT |
| Rab20 | TRCN000010264 0 | CCTGACAGAAACAGCCAACAA |
| Rab20 | TRCN000010264 2 | CCTCCTCTTTGAAACCTTGTT |
| Rab20 | TRCN000010264 3 | CCCTTTACAAGAAGATCCTGA |
| Rab20 | TRCN000010264 4 | GAAGATCCTGAAGTACAAGAT |
| Rab21 | TRCN000010268 0 | CCTACTTAAAGGCTTCATTTA |
| Rab21 | TRCN000010268 1 | GCCCAATTTACTACCGAGATT |
| Rab21 | TRCN000010268 2 | GCATTACCATACTTCCGCTAA |
| Rab21 | TRCN000010268 3 | GCAGGCATCTTTCTTAACAAA |
| Rab21 | TRCN000010268 4 | GCTAAACAGAACAAAGGCATT |

| | | |
|---|---|---|
| Rab22a | TRCN0000100830 | GCCGGTGTGTATTGATCGTTT |
| Rab22a | TRCN0000100833 | GCTGGACAAGAACGATTTCGT |
| Rab22a | TRCN0000100834 | CGCAGATTCCATTCATGCCAT |
| Rab22a | TRCN0000302790 | GCAGGGAACAAGTGCGATCTT |
| Rab22a | TRCN0000302791 | CCGAATATCAATCCAACCATA |
| Rab22a | TRCN0000376917 | ATGGATGGTAGGATTGAATTG |
| Rab22a | TRCN0000380076 | GACGCCACCTCATGCTCTTTA |
| Rab22a | TRCN0000381994 | TGTCAGAGTCGTATCAGTAAG |
| Rab23 | TRCN0000102770 | GCCAAACACTACCGCTAATTT |
| Rab23 | TRCN0000102771 | GCTGGATGATTCATGCATCAA |
| Rab23 | TRCN0000102772 | CGACAGATTCAGGTTAACGAT |
| Rab23 | TRCN0000102773 | ACACTCAAGTAGTAACAAGAT |
| Rab23 | TRCN0000102774 | CATCAACCTTAGACCTAACAA |
| Rab24 | TRCN0000102805 | CCCAGTGGAATTAGATGAATT |
| Rab24 | TRCN0000102806 | CGTCGTGTAGACTTCCATGAT |
| Rab24 | TRCN0000102807 | GCCGATAATATCAAAGCCCAA |
| Rab24 | TRCN0000102809 | AGCCATGAGCAGAATCTATTA |
| Rab24 | TRCN0000306308 | ATCTACCTGTGTGGCACTAAG |
| Rab24 | TRCN0000326454 | GAGAGCCAAGTTCTGGGTTAA |
| Rab24 | TRCN0000382512 | TCAGGACTATGCCGATAATAT |
| Rab25 | TRCN0000100250 | CCTGCCTTCAGCTTTCAGATA |
| Rab25 | TRCN0000100251 | CCTGGTATTTGACCTGACCAA |
| Rab25 | TRCN0000100252 | GCCCTCGACTCCACCAATGTT |
| Rab25 | TRCN0000100253 | CTTTGTCTTTAAGGTGGTGCT |
| Rab25 | TRCN000010025 | CCTCAAAGAGATCTTTGCAAA |

| | | |
|---|---|---|
| | 4 | |
| Rab25 | TRCN0000238206 | ATATCTCCACCTCCCTTACTG |
| Rab25 | TRCN0000238207 | GCTGGCTAAAGGAGCTGTATG |
| Rab25 | TRCN0000238208 | CATGCCGAAGCCACGATTGTT |
| Rab26 | TRCN0000341409 | TGAGAATTGCCCGAGTCTATA |
| Rab26 | TRCN0000341411 | CAGGCTGCATGACTATGTTAA |
| Rab26 | TRCN0000341412 | TCTACGACATCACCAACAAAG |
| Rab26 | TRCN0000341413 | ACTCAAGACCGTGTGGTAAAG |
| Rab26 | TRCN0000341481 | GGCATCGACTTCCGGAATAAA |
| Rab27a | TRCN0000100575 | GCCAGTTTAAGAGAAGTGTTT |
| Rab27a | TRCN0000100576 | CGAAACTGGATAAGCCAGCTA |
| Rab27a | TRCN0000100577 | GCTTCTGTTCGACCTGACAAA |
| Rab27a | TRCN0000100578 | CCAGTACACTGATGGCAAGTT |
| Rab27a | TRCN0000100579 | GACAAACATAAGCCACGCGAT |
| Rab27b | TRCN0000100425 | CCTGAGACAATGTCAAACCAT |
| Rab27b | TRCN0000100426 | CGGGAAGACAACATTTCTCTA |
| Rab27b | TRCN0000100427 | GCATACCATACTTCGAAACAA |
| Rab27b | TRCN0000100428 | GCTTCTGGACTTAATCATGAA |
| Rab27b | TRCN0000100429 | CTCTATAGATACACAGACAAT |
| Rab28 | TRCN0000100695 | GCTTCAAATGAGGCTGTAATA |
| Rab28 | TRCN0000100696 | CCTGGGAATCAAATTAAACAA |
| Rab28 | TRCN0000100697 | GCAGACGATAGGACTGGATTT |
| Rab28 | TRCN0000100698 | GCACACACTGTATTGATAGTT |
| Rab28 | TRCN0000100699 | GCTGATAAGCACTTACGATTT |
| Rab7l1 | TRCN0000102730 | CTCTGAGTCATTCCAATTCTT |

| | | |
|---|---|---|
| Rab7l1 | TRCN0000102731 | CCATGACACGACTCTACTATA |
| Rab7l1 | TRCN0000102732 | GTTCAGTAAAGAGAATGGCTT |
| Rab7l1 | TRCN0000102733 | GACTCTACTATAGAGATGCTT |
| Rab7l1 | TRCN0000102734 | CAATGCCACTACTTTCAGCAA |
| Rab30 | TRCN0000100401 | CAGCTATTTGACTTGTTGTAA |
| Rab30 | TRCN0000100402 | CGAAGATTCACTCAGGGTCTT |
| Rab30 | TRCN0000348722 | ACGTGCCTAGTCCGAAGATTC |
| Rab30 | TRCN0000351883 | CGGGAGATAGAACAGTATGCT |
| Rab30 | TRCN0000351884 | GCAGTCTTCTTCAGTCTCATT |
| Rab30 | TRCN0000351959 | CTTGATCCTTACCTATGACAT |
| Rab31 | TRCN0000100435 | GCGGGTTTGTATATGTGTAAA |
| Rab31 | TRCN0000100436 | GCAGGATTCATTTCATACCTT |
| Rab31 | TRCN0000100437 | CCAGGATCACTTTGACCACAA |
| Rab31 | TRCN0000100438 | CAGCGCGAAGAATGCCATTAA |
| Rab31 | TRCN0000100439 | CGCGAAGAATGCCATTAACAT |
| Rab31 | TRCN0000379445 | ACCGTGCCTTGTGGAAATGAA |
| Rab31 | TRCN0000380370 | GGGTTGGGAAATCCAGCATTG |
| Rab31 | TRCN0000380682 | CGCTAAGGAGTACGCTGAATC |
| Rab31 | TRCN0000381460 | TCAAGGAATCAGCCGCCAGAT |
| Rab31 | TRCN0000381947 | TGGCGATTGCTGGGAACAAGT |
| Rab32 | TRCN0000102686 | CTACATTTGATGCAGTCCTAA |
| Rab32 | TRCN0000102687 | CCTCTGCCAAGGATAATATAA |
| Rab32 | TRCN0000102688 | CGTGGGTAAGACGAGCATCAT |
| Rab32 | TRCN0000102689 | GAAATCGACCTGGACAGAATT |
| Rab32 | TRCN000028845 | GCCAAGTTTCTGTAGTGTAAA |

| | | |
|---|---|---|
| | 0 | |
| Rab33a | TRCN000010072 5 | GCTCATAACATGCTCTTGTTT |
| Rab33a | TRCN000010072 6 | GACCTCCTTCACCAACTTAAA |
| Rab33a | TRCN000010072 7 | CTCCAACTTAGCCCTGAAATT |
| Rab33a | TRCN000010072 8 | CAGTACGTGCAGATTCGCATT |
| Rab33b | TRCN000010060 0 | GCCCAATACATCTTTCTTAAT |
| Rab33b | TRCN000010060 1 | GCGAACGACATACCTCGAATT |
| Rab33b | TRCN000010060 2 | GATAACAGAATTAGCCTGAAA |
| Rab33b | TRCN000010060 3 | GCTTGGATAGAGGAATGCAAA |
| Rab33b | TRCN000010060 4 | CCACGTAGAAGCTATATTCAT |
| Rab34 | TRCN000010052 0 | CCTGGACATTTGCACTGACTT |
| Rab34 | TRCN000010052 1 | CCAAGAGATTAAGGCCGAGTA |
| Rab34 | TRCN000010052 2 | GCACTAACCTTTGAGGCCAAT |
| Rab34 | TRCN000010052 3 | CGTGGGATTTAAGATATCCAA |
| Rab34 | TRCN000030268 6 | CGGCACATTGCAGATGTTGTT |
| Rab34 | TRCN000030493 0 | CTGCAAAGACACCTTCGATAA |
| Rab34 | TRCN000030493 2 | TGAGTACTCCTGCTCAGTATT |
| Rab34 | TRCN000031116 8 | GGCTACCATCGGAGTGGATTT |
| Rab34 | TRCN000037443 2 | ATCGTTGTGGGAGACCTATCT |
| Rab34 | TRCN000038086 9 | GACAGGCACCGTGGGATTTAA |
| Rab35 | TRCN000010053 0 | CGTAACTCAGAAGAACTGATT |
| Rab35 | TRCN000010053 1 | GCCGAATATTAGTGGGCAATA |
| Rab35 | TRCN000010053 2 | GCTGTTACGATTCGCAGACAA |
| Rab35 | TRCN000010053 3 | CGAGTCCTTTGTCAACGTCAA |
| Rab35 | TRCN000010053 4 | AGTGCCAAGGAGAACGTCAAT |

| | | |
|---|---|---|
| Rab35 | TRCN0000379544 | CTCAGTTTAGTGCCGTTATTT |
| Rab35 | TRCN0000379586 | CAAGCGATGGCTTCATGAAAT |
| Rab35 | TRCN0000379707 | ACCATCACCTCTACGTATTAT |
| Rab35 | TRCN0000380231 | GACAGAAGATGCCTACAAATT |
| Rab36 | TRCN0000100800 | CCCATCTTTCTCCTGCCATAA |
| Rab36 | TRCN0000100801 | CCATGATTACAAGGCCACGAT |
| Rab36 | TRCN0000100802 | CCTCATTCACAGGTTGTGCAA |
| Rab36 | TRCN0000100803 | GATGGAGACCTAATACGAATA |
| Rab36 | TRCN0000100804 | CAAGTGTATTGCGTCTGCCTA |
| Rab37 | TRCN0000100815 | GCAGAACTGAACAAAGCCATA |
| Rab37 | TRCN0000100816 | CCAGGGAATATGGTGTTCCTT |
| Rab37 | TRCN0000100817 | GACGTGGTGATTATGCTTCTA |
| Rab37 | TRCN0000100818 | GCCCAGAGAGACGTGGTGATT |
| Rab37 | TRCN0000100819 | CGGCATAGACTCCAGGAATAA |
| Rab38 | TRCN0000102645 | GCCCTAATATTTGTTCCTTTA |
| Rab38 | TRCN0000102646 | GCTTCGTAGGATGGTTTGAAA |
| Rab38 | TRCN0000102647 | GAGTCTATAGAACCGGACATT |
| Rab38 | TRCN0000102648 | ACCAGCATTATCAAGCGCTAT |
| Rab38 | TRCN0000102649 | CACATTTGAAGCCGTGGCAAA |
| Rab39 | TRCN0000102695 | CCGTCTTAGAAATACTGACTA |
| Rab39 | TRCN0000102696 | CGCAACTCAGTTGGAGGATTT |
| Rab39 | TRCN0000102697 | GCTTCAGATCAATAACTCGAT |
| Rab39 | TRCN0000102698 | AGATCAATAACTCGATCCTAT |
| Rab39 | TRCN0000102699 | GACTGTGGAATGAAGTACATA |
| Rab39b | TRCN000010274 | CCCAGGATTATCCAGTGGATA |

| | | |
|---|---|---|
| | 5 | |
| Rab39b | TRCN000010274 6 | CCCTACCAAATTGTATTTGTT |
| Rab39b | TRCN000010274 7 | CGGCTCATTGTCATCGGCGAT |
| Rab39b | TRCN000010274 8 | CGCTTTGCTCAGGTTTCAGAT |
| Rab39b | TRCN000010274 9 | GCCTACTACAGGAATTCAGTA |
| Rab40b | TRCN000010276 5 | GCATTTATTAACGGTAACGAT |
| Rab40b | TRCN000010276 6 | CGATGGATTAAGGAGATTGAT |
| Rab40b | TRCN000010276 7 | CCACCTTAAGTCTTTCTCGAT |
| Rab40b | TRCN000010276 8 | CTTCAACATTACAGAGTCCTT |
| Rab40b | TRCN000010276 9 | GTGGTCCTGGTCTATGACATT |
| Rab40c | TRCN000005484 9 | GCCGTGCCATTGTCTCCTGTA |
| Rab40c | TRCN000005485 0 | CTGTACCATCTTCAGGTCCTA |
| Rab40c | TRCN000005485 1 | GCAACAGCCTTAAGAGGTCTA |
| Rab40c | TRCN000005485 2 | CCACTACCTGTCACCATCAAA |
| Rab40c | TRCN000028739 2 | CTGTGCAACTTCAACGTCATT |
| Rab40c | TRCN000029488 7 | AGCAACGGGATAGACTATAAG |
| Rab40c | TRCN000030744 9 | AGTGTTGAAGCCAGATCTTTA |
| Rab40c | TRCN000038051 3 | TGGATCAAGGAGATCGATGAG |
| Rab42 | TRCN000034146 0 | ACGTCACTGCTACGGTGTTAC |
| Rab42 | TRCN000034146 4 | CTGACAAGGTGGTCTTCTTAC |
| Rab42 | TRCN000034146 6 | TGACGAACAGAGAGTCCTTTG |
| Rab42 | TRCN000034146 7 | TGGGCGTTCTGTTGGTCTTTG |
| Rab42 | TRCN000034146 9 | ACCTGAACACCCGATGCGTAT |
| Rab43 | TRCN000010281 1 | AGGTCCCATGTTCAGTGAGAA |
| Rab43 | TRCN000031824 8 | ACAAGTCAGACCTTGCCGATT |

| | | |
|---|---|---|
| Rab43 | TRCN0000318249 | GAGCACTATGACATCCTCTGT |
| Rab43 | TRCN0000318322 | GATCGAGGATGTGAGGAAGTA |
| Rab43 | TRCN0000318323 | CCTGCCTCTTACCAAAGTATA |
| Rabl2a | TRCN0000100885 | CCCTACATTTCTGGAGAACAT |
| Rabl2a | TRCN0000100886 | GCAGACATACAGATGACTCAA |
| Rabl2a | TRCN0000100887 | CCCAAGACACAGACCCATATT |
| Rabl2a | TRCN0000100888 | GCACCACACATTTCCTTCTTT |
| Rabl2a | TRCN0000100889 | GTCCAAACTCATGGAGAGATT |
| Rabl3 | TRCN0000181097 | CCCAGAGTAACGAACTCATAT |
| Rabl3 | TRCN0000182964 | CACAATCAAGTGTTAGGAAAT |
| Rabl3 | TRCN0000241041 | TGAAGAGAAGACATACTATAT |
| Rabl3 | TRCN0000241042 | TTGCTGCTGAAGGCTTAATTA |
| Rabl4 | TRCN0000102720 | ACTGTCCTGAAGGGCTGCCCA |
| Rabl4 | TRCN0000102721 | GCCTGGAATTCTTTGAGACAT |
| Rabl4 | TRCN0000102722 | CCTTGTCTATGATGTGACCAA |
| Rabl4 | TRCN0000102723 | CTGGATAAGTTGTGGGAGAAT |
| Rabl4 | TRCN0000102724 | CAGTGCCAGTTCTTGACACAA |
| Rabl5 | TRCN0000191434 | CCACCTAAAGGAAATTGAAAT |
| Rabl5 | TRCN0000191988 | GTGGAGTTCATCAAGTATTTA |
| Rabl5 | TRCN0000200816 | GCCACCTAAAGGAAATTGAAA |
| Rabl5 | TRCN0000215745 | GATCGTCTTCAATGCTGATAT |
| Rabl5 | TRCN0000247292 | GGTGGAGTTCATCAAGTATTT |
| Rabl5 | TRCN0000247293 | TCCCAAAGGACGCTGACTAAC |
| Rabl5 | TRCN0000257636 | TCGTCTTCAATGCTGATATTC |
| Rabl5 | TRCN000025765 | CTGAAGCTGGTGCACTCAAAC |

| | | |
|---|---|---|
| | 7 | |
| Rabl5 | TRCN000025776<br>2 | CTTCGGACATCACTGAATATA |
| Hps-1 | TRCN000029255<br>6 | GCAAGCTGTTGGCTTTCTACT |

**Table S4.** Candidate genes identified from the screen and shRNA for validation

**(A)** List of candidate genes identified from the screen

| Z-score (>1.18) | Gene name | TRCN |
|---|---|---|
| 2.83 | Rab37 | TRCN0000100817 |
| 2.70 | Rab25 | TRCN0000100252 |
| 2.68 | Rab35 | TRCN0000379586 |
| 2.60 | Rab35 | TRCN0000100532 |
| 2.46 | Rab27a | TRCN0000100577 |
| 2.38 | Rab22a | TRCN0000302790 |
| 2.35 | Rab40b | TRCN0000102769 |
| 2.29 | Rab32 | TRCN0000102688 |
| 2.28 | Rab43 | TRCN0000318248 |
| 2.13 | Rab40c | TRCN0000380513 |
| 2.13 | Rab31 | TRCN0000381460 |
| 2.07 | Rab25 | TRCN0000238207 |
| 1.99 | Rab24 | TRCN0000306308 |
| 1.86 | Rab37 | TRCN0000100818 |
| 1.83 | Rab10 | TRCN0000335543 |
| 1.78 | Rab1b | TRCN0000302710 |
| 1.77 | Rab28 | TRCN0000100697 |
| 1.75 | Rab30 | TRCN0000100402 |
| 1.74 | Rab11a | TRCN0000305864 |
| 1.70 | Rab1b | TRCN0000302711 |
| 1.69 | Rab1b | TRCN0000381762 |
| 1.65 | Rab36 | TRCN0000100804 |
| 1.65 | Rab31 | TRCN0000380682 |
| 1.65 | Rab7 | TRCN0000100881 |
| 1.63 | Rabl5 | TRCN0000257657 |
| 1.63 | Rab3c | TRCN0000089456 |
| 1.62 | Rab17 | TRCN0000100918 |
| 1.54 | Rab43 | TRCN0000318322 |
| 1.52 | Rab40c | TRCN0000054851 |
| 1.51 | Rab10 | TRCN0000335544 |
| 1.49 | Rab7 | TRCN0000100880 |
| 1.36 | Rab25 | TRCN0000100251 |
| 1.36 | Rab38 | TRCN0000102649 |
| 1.34 | Rab31 | TRCN0000100438 |
| 1.25 | Rab27a | TRCN0000100579 |
| 1.23 | Rab20 | TRCN0000102640 |
| 1.22 | Hps-1 | TRCN0000292556 |
| 1.20 | Rab15 | TRCN0000093173 |
| 1.18 | Rab32 | TRCN0000288450 |

**(B)** List of shRNA for validation

| **List of candidate genes/shRNA to validate** | | |
|---|---|---|
| **Z- score** | **Gene** | **TRCN** |
| 2.60 | Rab35 | TRCN0000100532 |
| 2.07 | Rab25 | TRCN0000238207 |
| 1.86 | Rab37 | TRCN0000100818 |
| 1.83 | Rab10 | TRCN0000335543 |
| 1.69 | Rab1b | TRCN0000381762 |
| 1.65 | Rab31 | TRCN0000380682 |
| 1.54 | Rab43 | TRCN0000318322 |
| 1.52 | Rab40c | TRCN0000054851 |
| 1.49 | Rab7 | TRCN0000100880 |
| 1.25 | Rab27a | TRCN0000100579 |
| **1.22** | **Hps-1** | **TRCN0000292556** |
| **1.18** | **Rab32** | **TRCN0000288450** |

**Table S5**. Primers used in this study

| NAME | SEQUENCE (5´-3´) |
|---|---|
| F-NGS | TCGTCGGCAGCGTCAGATGTGTATAAGAGACAGTCTTGTGGAAAGGACGA |
| R-NGS | GTCTCGTGGGCTCGGAGATGTGTATAAGAGACAGTCTACTATTCTTTCCCCTGC ACTGT |
| F-Hps1 | TGGTTCGAGAATGACATGGGA |
| R-Hps1 | GGGTGGCTCTTGCTGTAGTAG |
| F-Rab1b | ATGAACCCCGAATATGACTACCT |
| R-Rab1b | TGCTGATGTAGCTCTCTGTGTA |
| F-Rab7 | AAGCCACAATAGGAGCGGAC |
| R-Rab7 | AGACTGGAACCGTTCTTGACC |
| F-Rab10 | GGCAAGACCTGCGTCCTTTT |
| R-Rab10 | GTGATGGTGTGAAATCGCTCC |
| F-Rab25 | GGGTTGAGTTCTCCACCCG |
| R-Rab25 | CCCCACGATAGTACGCAGA |
| F-Rab27a | TCGGATGGAGATTACGATTACCT |
| R-Rab27a | TTTTCCCTGAAATCAATGCCCA |
| F-Rab31 | GACACGGGGGTTGGGAAATC |
| R-Rab31 | ACAAGGCACGGTTTTGGTCA |
| F-Rab32 | CGTGGGTAAGACGAGCATCAT |
| R-Rab32 | CCCAGTTGAGAACTTTGAGGG |
| F-Rab35 | CCACAATCGGAGTGGATTTCA |
| R-Rab35 | CGTCGTAAACCACAATGACCC |
| F-Rab37 | CCAACCAGTCCTCTTTTGACAA |
| R-Rab37 | GCCTAGAAGCATAATCACCACG |
| F-Rab40c | GGGCAATAAGAATGATGACCCTG |
| R-Rab40c | TCCACATTGACGTTCTCCTTG |
| F-Rab43 | CAAGCTGGTGTTAGTGGGC |
| R-Rab43 | CCCAAATCTGTAACTTGACCCG |
| F-GAPDH | AGGTCGGTGTGAACGGATTTG |
| R-GAPDH | TGTAGACCATGTAGTTGAGGTCA |
